# Widespread PERK-dependent repression of ER targets in response to ER stress

**DOI:** 10.1101/487934

**Authors:** Nir Gonen, Niv Sabath, Christopher B. Burge, Reut Shalgi

## Abstract

The UPR (Unfolded Protein Response) is a well-orchestrated response to ER protein folding and processing overload, integrating both transcriptional and translational outputs. Its three arms in mammalian cells, the PERK translational response arm, together with the ATF6 and IRE1-XBP1-mediated transcriptional arms, have been thoroughly investigated.

Using ribosome footprint profiling, we performed a deep characterization of gene expression programs involved in the early and late ER stress responses, within WT or PERK -/- Mouse Embryonic Fibroblasts (MEFs). We found that both repression and activation gene expression programs, affecting hundreds of genes, are significantly hampered in the absence of PERK. Specifically, PERK -/- cells do not show global translational inhibition, nor do they specifically activate early gene expression programs upon short exposure to ER stress. Furthermore, while PERK -/- cells do activate/repress late ER-stress response genes, the response is substantially weaker. Importantly, we highlight a widespread PERK-dependent repression gene expression program, consisting of ER targeted proteins, including transmembrane proteins, glycoproteins, and proteins with disulfide bonds. This phenomenon occurs in various different cell types, and has a major translational regulatory component. Moreover, we revealed a novel interplay between PERK and the XBP1-ATF6 arms of the UPR, whereby PERK attenuates the expression of a specific subset of XBP1-ATF6 targets, further illuminating the complexity of the integrated ER stress response.

## Introduction

Protein homeostasis is one of the hallmarks of cellular viability and a well-known factor in health and disease. Rapidly changing cellular environments demand robust cellular and molecular responses, enabling cell survival under extreme conditions. The endoplasmic reticulum (ER) is a main regulator for cellular protein homeostasis, translating up to 50% of all proteins in certain cells^1^. The ER is a hub for translation and trafficking of membrane bound, integrated membrane, and secreted proteins^2,3^. Furthermore, numerous proteins are subject to major post-translational modifications inside the ER, including disulfide bond formation and glycosylation^3^. ER-stress has long been known to elicit a complex cellular program, also termed the Unfolded Protein Response (UPR), which has evolved to allow cells to cope with dynamic changes in the protein folding and processing demands in the ER^2,4,5^. The metazoan UPR consists of three evolutionary distinct branches: IRE1-XBP1, ATF6 and the protein kinase RNA-like endoplasmic reticulum kinase (PERK)^2,6^. While IRE1-XBP1 and ATF6 are known to mediate a transcriptional response, the PERK arm primarily elicits a global translational response, with a secondary, ATF4-mediated transcriptional component^7^. PERK has been shown to phosphorylate the Eukaryotic Initiation Factor 2 α (eIF2α) translation initiation factor, thereby inhibiting ribosomal ternary complex recycling^7,8^, to reduces global translation initiation rates. The secondary ATF4-dependnet transcriptional response induces a variety of genes necessary for adaptation to ER overload^2^. Accordingly, ATF4 upregulates the GADD34 phosphatase, which leads to eIF2α dephosphorylation, and subsequent relaxation in the translation initiation repression^2^.

Recent work has made a distinction between acute, early ER-stress response and chronic ER-stress, which is considered most relevant to disease^5,9^, occurring at the stage of eIF2α-phosphorylation relief and partial translational relaxation. Furthermore, a major role for eIF3-dependent translation during the chronic stage was described^9^. Additionally, a transient shift in the localization of mRNAs encoding membrane and secreted proteins away from ER-bound ribosomes towards cytosolic ribosomes has been reported to ensue shortly after triggering ER stress^10^.

PERK knockout (PERK -/-) cells have been useful for establishing PERK’s function in cellular homeostasis maintenance under ER-stress^11^. Previous genome-wide studies have used mRNA expression profiling to define a transcriptional response following a 6h ER-stress in PERK -/- and ATF4 -/- cells^12,13^. These experiments have shown PERK-dependent metabolic changes enabling the maintenance of redox potential under ER-stress^13^.

Continuing the wide body of research on the role of PERK in ER stress, we sought to understand the early and sustained PERK-dependent components of the UPR in a transcriptome-wide manner. While the translational arm of the UPR is fairly immediate, the impact of the transcriptional arms on cellular gene expression takes time to manifest. Thus, the different arms of the UPR generate a complex integrated regulation of gene expression programs in various stages of the response. Furthermore, while PERK is known to elicit an eIF2α phosphorylation-mediated global translational repression in response to ER stress, its role in controlling the translation of specific gene expression programs still remains elusive. We therefore chose to approach these questions in a manner that examines gene expression programs as an integration of transcription and translation.

In this study we examined the PERK-dependent dynamic alterations in gene expression programs following ER-stress using ribosome footprint profiling^14^ on Wild-Type (WT) and PERK -/- Mouse Embryonic Fibroblasts (MEFs)^11^ treated with ER stress. We used Thapsigargin (Tg), a SERCA inhibitor, for short (1h or 2h) or long (5h or 8h) treatments, to examine the gene expression programs that govern either early or late responses to ER stress.

We characterized three major gene expression programs in response to ER stress in PERK WT MEFs: Early induction, late induction and repression. We further show that all three programs are markedly compromised in the absence of PERK. We describe a widespread, PERK-dependent repression program consisting of ER target genes, including transmembrane proteins, glycoproteins, and proteins with disulfide bonds. Finally, we reveal that PERK attenuates a specific subset of XBP1-ATF6 target genes, thereby unraveling a complex interplay between the different arms of the UPR.

## Results

### Exploring PERK dependent early and late ER stress gene expression programs

In order to identify gene expression programs that are activated or repressed in a PERK dependent manner during ER stress, we used Mouse Embryonic Fibroblast (MEF) cells that are either WT or were knocked out for PERK (PERK -/-)^11^. We treated these cells with Tg (1 µM), for various durations to assay both the early, immediate, response to ER stress (1h or 2h), as well as the late adaptive ER stress response (5h or 8h).

We first performed polysome profiling, to look at the global kinetics of the early and late translational responses ER stress. WT MEFs exhibited marked translational repression, which was maximal at 1h of ER stress, and showed a time dependent relaxation, indicative of gradual adaptation (Fig. 1A). In contrast, PERK -/- MEFs showed no change in their global overall translation levels, and only at the 8h timepoint a slight repression of translation was observed (Fig. 1B). However, even at 8h, the level of translational repression observed in PERK -/- MEFs did not resemble nearly any of the treatments of WT MEFs.

**Figure 1.**
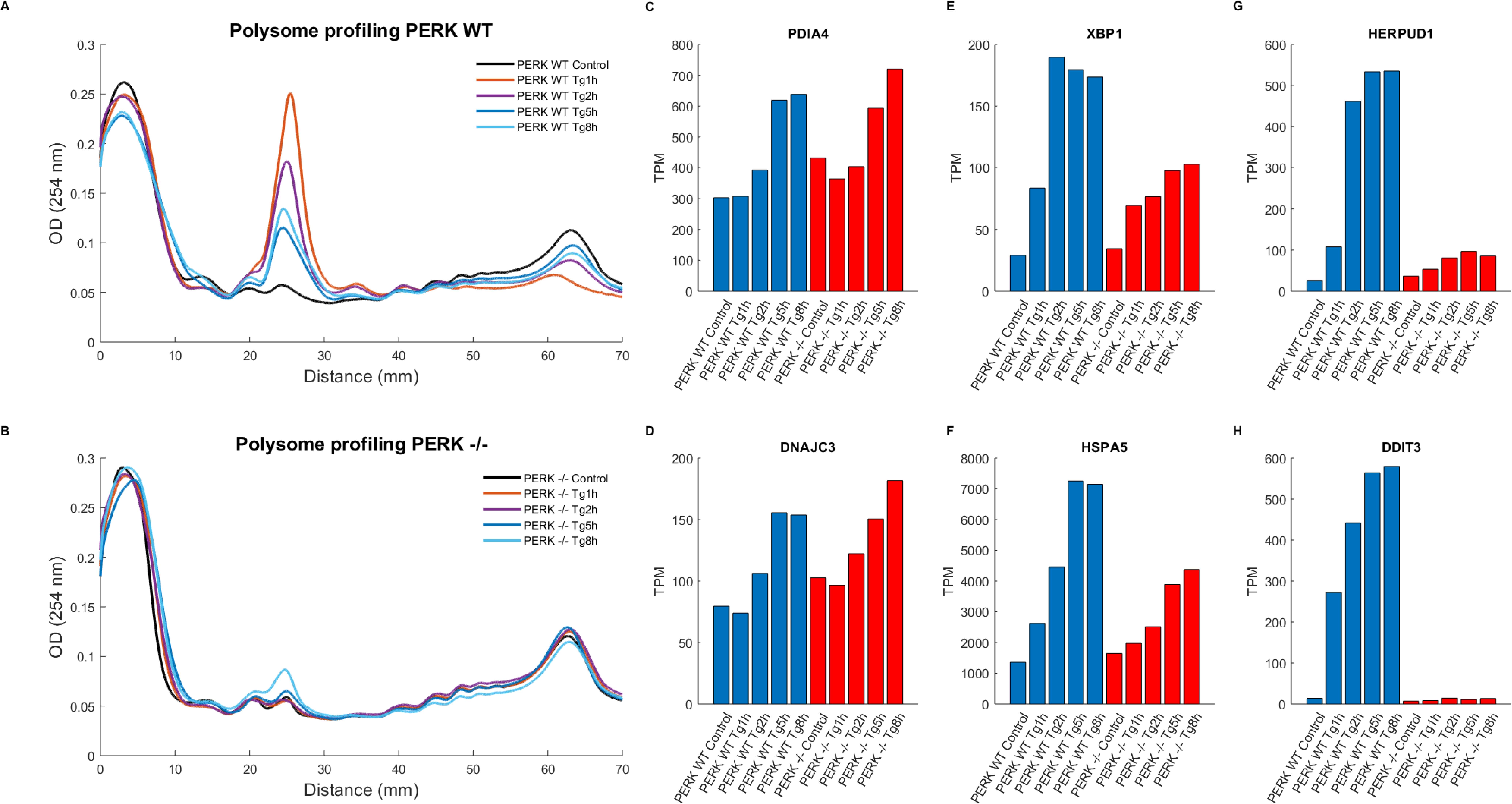
PERK-dependent translational repression dynamics. (A,B) Polysome profiling demonstrated overall translational repression and partial recovery upon ER stress in PERK WT MEFs but not in PERK -/- MEFs. RNAs were separated on a sucrose gradient (10%-50%) using an ultracentrifuge, and the gradient was read using a UV reader. The graph shows the amount of RNA bound by different size polysomes in the different conditions. Polysome/monosme ratios (P/M), used to measure the level of overall translation, as the ratio between the polysome area under the curve (4- somes and up) to the monosome area under the curve, were calculated throughout Tg treatment in PERK WT and PERK -/- cells. (A) PERK WT P/M decreased sharply following Tg treatment and showed relaxation in a time-dependent manner: Control P/M = 5.97; Tg 1h P/M = 1.17; Tg 2h P/M: 1.79; Tg 5h P/M =2.85; Tg 8h P/M = 2.38. (B) PERK -/- P/M did not show an immediate decrease following Tg treatment, with the exception of a slight repression after 8h: Control P/M= 6.35; Tg 1h P/M **=** 6.44; Tg 2h P/M =6.55; Tg 5h = 5.72; Tg 8h= 4.22. (C-H) Selected ER stress response genes altered during Tg treatment in PERK WT and PERK -/- cells. All bar plots depict gene expression TPM values in PERK WT and PERK -/- cells following Tg treatments (Control, Tg 1h, Tg 2h, Tg 5h, Tg 8h). ER-stress related genes showed three distinct activation patterns: while PDIA4 and DNAJC3 were activated in an PERK-independent manner (C,D); XBP1 and HSPA5 (BiP) have shown a partial dependence on PERK (E, F); HERPUD1 and DDIT3 (CHOP) showed a complete dependence on PERK (G, H).

We then performed ribosome footprint profiling, to look at the gene expression programs that are regulated under the different conditions. Ribosome footprint profiling provides relative protein synthesis levels in each sample, allowing us to characterize mRNAs which translation is preferentially enhanced or repressed (see Materials and Methods).

Principle Component Analysis (PCA) of the data showed a clear separation between WT and PERK -/- MEFs, as well as a gradual temporal separation of ER stress durations (Fig. S1A). Classical ER stress target genes showed one of three expression patterns: an ER stress-mediated PERK independent upregulation (Fig. 1C,D), an ER stress-mediated upregulation that was partially PERK-dependent, such as in the cases of BiP (HSPA5) and XBP1 (Fig. 1E,F), or complete PERK dependence, with the examples of CHOP (DDIT3) and HERPUD1 (Fig. 1G,H). Finally, ATF4 translation showed the expected pattern of uORF (upstream Open Reading Frame) translation^15^, with a marked temporal increase in ribosome occupancy on its main ORF following ER stress in WT MEFs (Fig. S1B). PERK -/- MEFs did not show any ATF4 translation prior to, or upon ER stress (Fig. S1B), consistent with the lack of eIF2α-phosphorylation in these cells upon ER stress^11^.

Interestingly, examining the positional occupancy of ribosome footprints along ORFs showed a weak but significant accumulation of ribosomes at the 5’ ends of ORFs, until around position 140 bases downstream to the AUG (Fig. S2A). This is consistent with a similar observation from a recent study^10^. We and others have previously observed ribosome accumulation at 5’ ends of ORFs in similar positions following heat shock^16^ and another proteotoxic stress condition^17^ and showed that they are consistent with ribosomes paused in elongation. In the case of ER stress, however, while highly significant (p-value<10^-300^ in all timepoints relative to control, using a KS-test), the magnitude of pausing was much smaller than observed under heat shock. Specifically, a ∼25-35% increase in pausing was observed under ER-stress at different times compared to control, compared to a 3-fold increase observed in heat shock. Notably, PERK -/- MEFs did not show any 5’-end ribosome accumulation (Fig. S2B).

### Ribosome footprint profiling revealed three major gene expression programs in response to ER stress

Next, we turned to ask what were the major gene expression programs that are characteristic of the early and late ER stress responses. To this end, we performed a clustering analysis of the Transcripts Per Million (TPM) values of the 1658 genes that changed their expression under ER stress in PERK WT cells (see Materials and Methods). The analysis revealed three major gene expression programs: (1) Early induction (Fig. 2, red cluster), containing 505 genes that were markedly upregulated in either 1 or 2h following stress, and somewhat relaxed during 5h and 8h of stress; (2) Late induction (Fig. 2, blue cluster), including 495 genes that gradually increased in their expression and showed a maximal induction at the 5h and 8h timepoints; (3) Repression (Fig. 2, black cluster), containing 658 downregulated genes.

**Figure 2.**
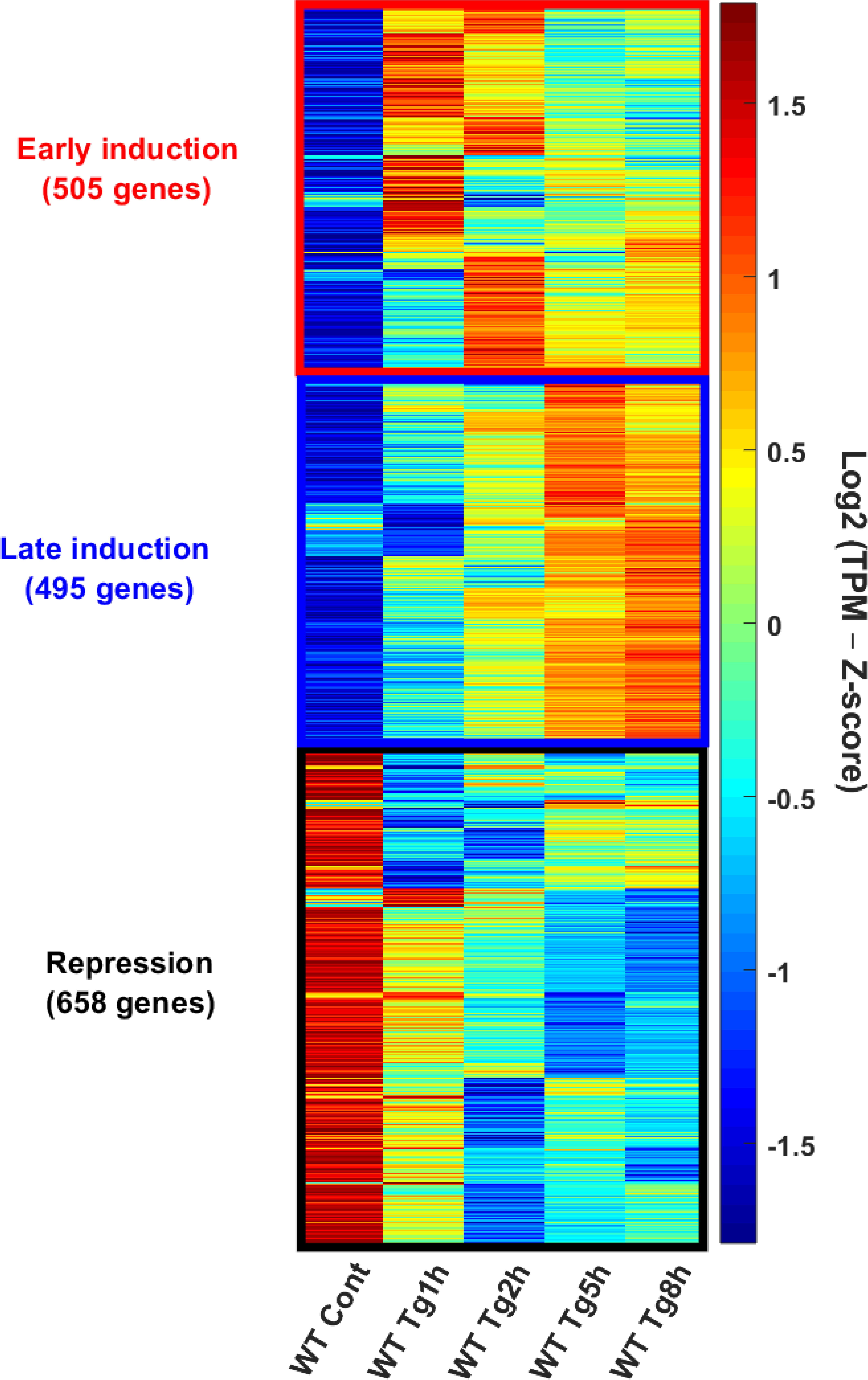
Major gene expression programs in response to ER stress. Hierarchical clustering analysis was performed using 1658 genes that were changed at least two-fold compared to Control in any of the time points following Tg treatment (1h, 2h,5h and 8h) in WT MEFs. Heatmap depicting hierarchal clustered genes TPMs, according to their spearman correlation. Z-score normalization was further performed for visualization purposes. The genes clustered into three distinct groups: Early induction (red cluster): 505 genes showing maximal activation during early time-points (1,2 hr) and relaxation in later timepoints (5,8 hr); Late Induction (blue cluster): 495 genes with a gradual activation throughout later timepoints (5,8 hours); Repression (black cluster): 658 genes strongly reduced during all timepoints.

We compared our gene expression programs to a recently published polysome-sequencing data by Guan et al. of MEFs subject to either 1h or 16h of Tg treatment^9^, and reassuringly, our results were recapitulated in this dataset (Fig. S3A-C). Furthermore, we compared the gene expression in our PERK -/- cells to MEFs treated with a PERK inhibitor following 16h of ER stress from Guan et al., and found that genes that were induced or repressed in PERK -/- cells at 8h of Tg were also significantly induced or repressed respectively at 16h Tg-stressed cells treated with a PERK inhibitor (Fig S3D).

To further substantiate the robustness of the major gene expression programs that we have identified in WT MEFs, we performed an additional experiment in NIH3T3 cells. NIH3T3 cells were exposed to ER stress (Tg) for either 2h or 7h, followed by polysome profiling and ribosome footprint profiling. Polysome profiling showed a similar trend of overall translational inhibition at 2h of ER stress, which relaxed at the 7h timepoint (Fig. 3A). Using ribosome footprint profiling, we found that the three major gene expression programs were largely recapitulated in NIH3T3 cells (Fig. 3B-D, S4), demonstrating the generality of our findings.

**Figure 3.**
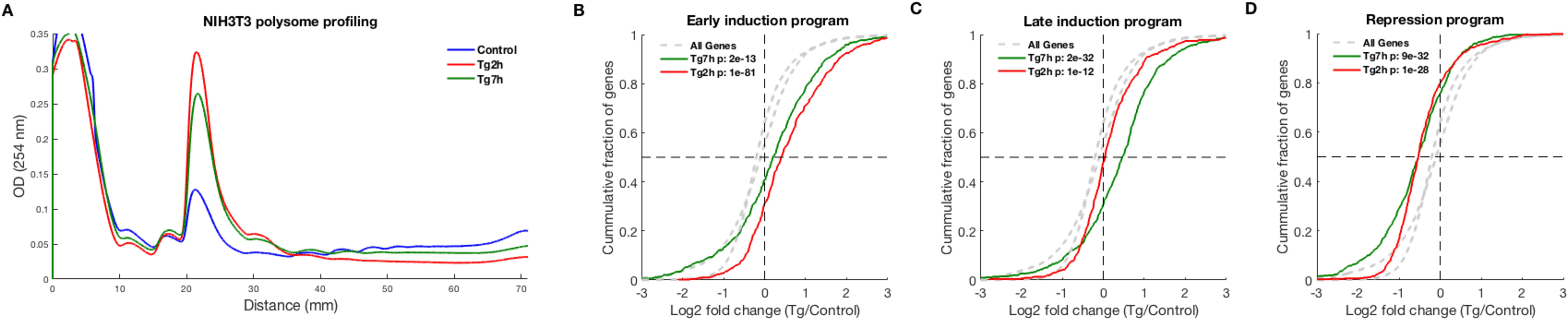
ER stress gene expression programs are recapitulated in NIH3T3 cells. (A) Polysome profiling of NIH3T3 cells shows major repression following 2h of Tg treatment, and a partial relaxation at 7h post Tg treatment (P/M ratios are: Control – 1.7, Tg 2h – 0.42, Tg 7h – 0.65). (B-D) Log2 fold changes (LFCs) of the genes from the three major ER stress gene expression programs, namely early induction (B), late induction (C) and repression (D), were calculated using ribosome footprint profiling data of Tg treated NIH3T3 cells for either 2h (red) or 7h (green). CDF plots demonstrate the cumulative fraction of each set of genes (y-axis), as a function of their log2 fold change (LFC) (x-axis) relative to control. Background distributions (LFC values of all expressed genes) in either 2h Tg or 7h Tg are marked in dashed grey lines. The analysis demonstrated that the three ER stress gene expression programs that were identified in MEFs were recapitulated in NIH3T3 cells. T-test p-values between each distribution and its respective background are indicated.

### Both early and late induction gene expression programs are PERK dependent

We next characterized which pathways are enriched within the induction gene expression programs. Functional enrichment analysis of the early induction program showed an enrichment for specific classes of histones as well as other DNA-binding proteins (Table S1A), as has been reported previously^10^. The late induction program was significantly enriched with genes annotated as “response to ER stress”, as expected (Table S1B). Additionally, the late induction program showed an enrichment for amino acid biosynthetic processes, in agreement with previous reports by Harding et al.^13^ using a different ER stressing agent, Tunicamycin (TM).

Examination of the early induction cluster showed that the increase in expression at the 1h and 2h timepoints is completely PERK dependent, while at the 5h and 8h timepoints, a very weak yet significant induction was observed in PERK -/- cells (Fig. 4A,B and S5A,B). Whereas the early induction program genes showed a median induction of 80% and 89% at the 5h and 8h of ER stress respectively, their median fold change was 12% and 11% in the PERK -/- cells at these timepoints.

**Figure 4.**
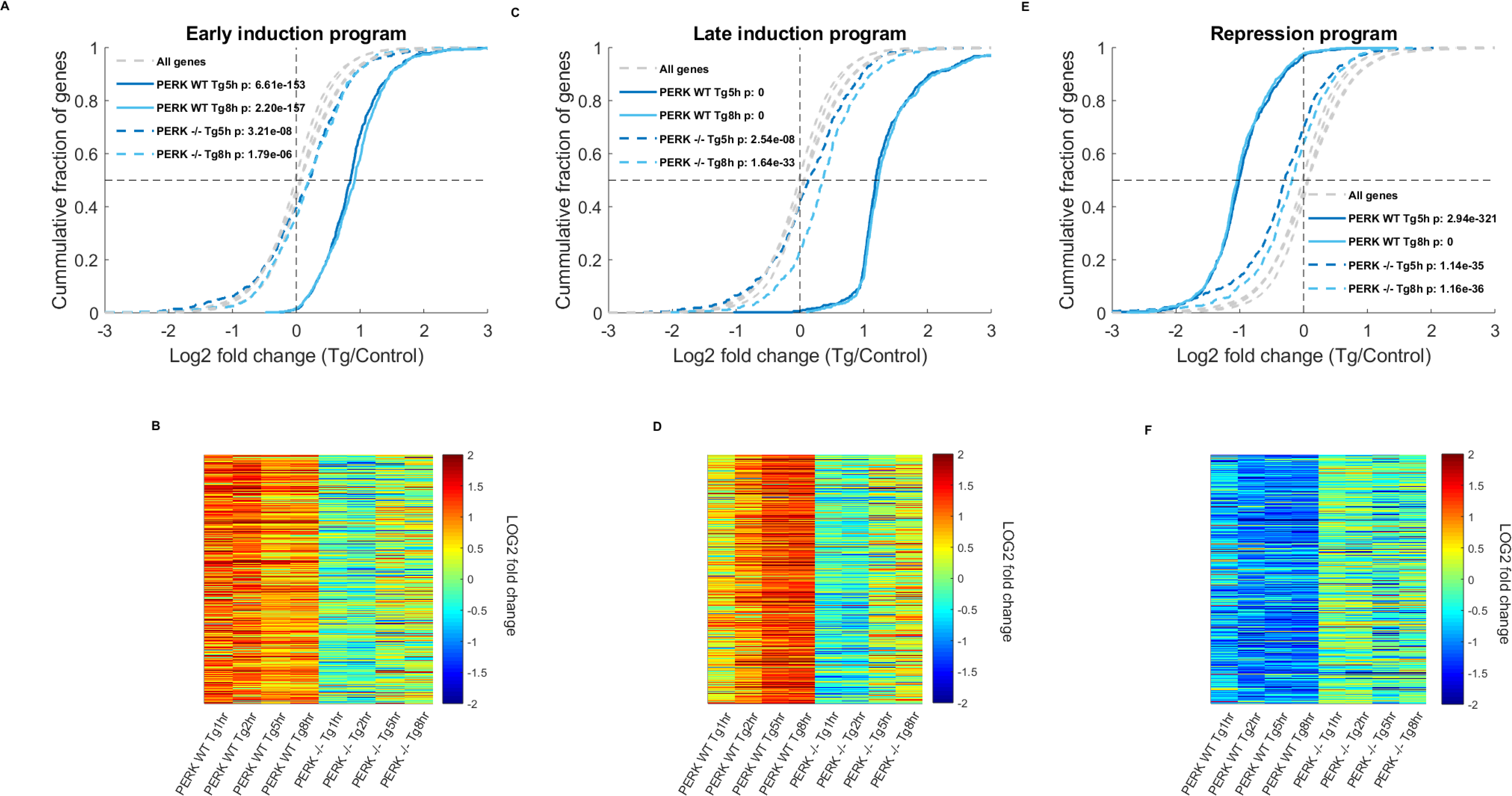
ER stress gene expression programs are impaired in the absence of PERK. (A,C,E) CDF plots depicting the log2 fold change (LFCs) of the early induction (A) late induction (C) and repression (E) gene expression programs within late timepoints. Grey dashed lines indicate the background distributions of the LFCs of all expressed genes in the different timepoints, t-test p-values between each distribution and its respective background are indicated. LFCs in PERK -/- cells (blue dashed lines) are significantly shifted, however the changes are extremely weak. (B,D,F) Heatmaps of LFCs of the early induction (B); late induction (D) and repression (F) gene expression program genes, clustered according to PERK WT TPMs, further illustrate the impaired regulation in PERK -/- cells.

Late induction cluster genes showed a slight aberrant repression in the early timepoints in PERK -/- MEFs (Fig. S5C,D), while at the later timepoints, a small yet significant induction was observed (Fig. 4C,D). Whereas the median fold change of the late induction genes was 128% and 136% in the PERK WT cells following 5h and 8h of Tg respectively, only 9% and 26% median induction values were observed in PERK -/- cells at these timepoints.

Taken together, we observed that PERK -/- cells were highly impaired in eliciting the desired gene expression program in the early response, (Fig. 4 and S5). Interestingly, at the late timepoints, cells were still unable to produce the full extent of activation of the required gene expression programs and showed a poor level of induction (Fig. 4A-D).

### PERK-mediated repression of ER targets during ER stress is a major component of the UPR

We next turned to characterize which pathways are enriched within the repression gene expression program. The repression gene expression program showed a significant enrichment for many functional annotation categories, which mainly converged upon two main themes (Fig. 5A, Table S1C): membrane, transmembrane, signal peptide-containing proteins and cell surface proteins, which represent a major class of ER-targeted proteins; and disulfide-bond containing proteins and glycoproteins, representing proteins undergoing post-translational modifications inside the ER.

**Figure 5.**
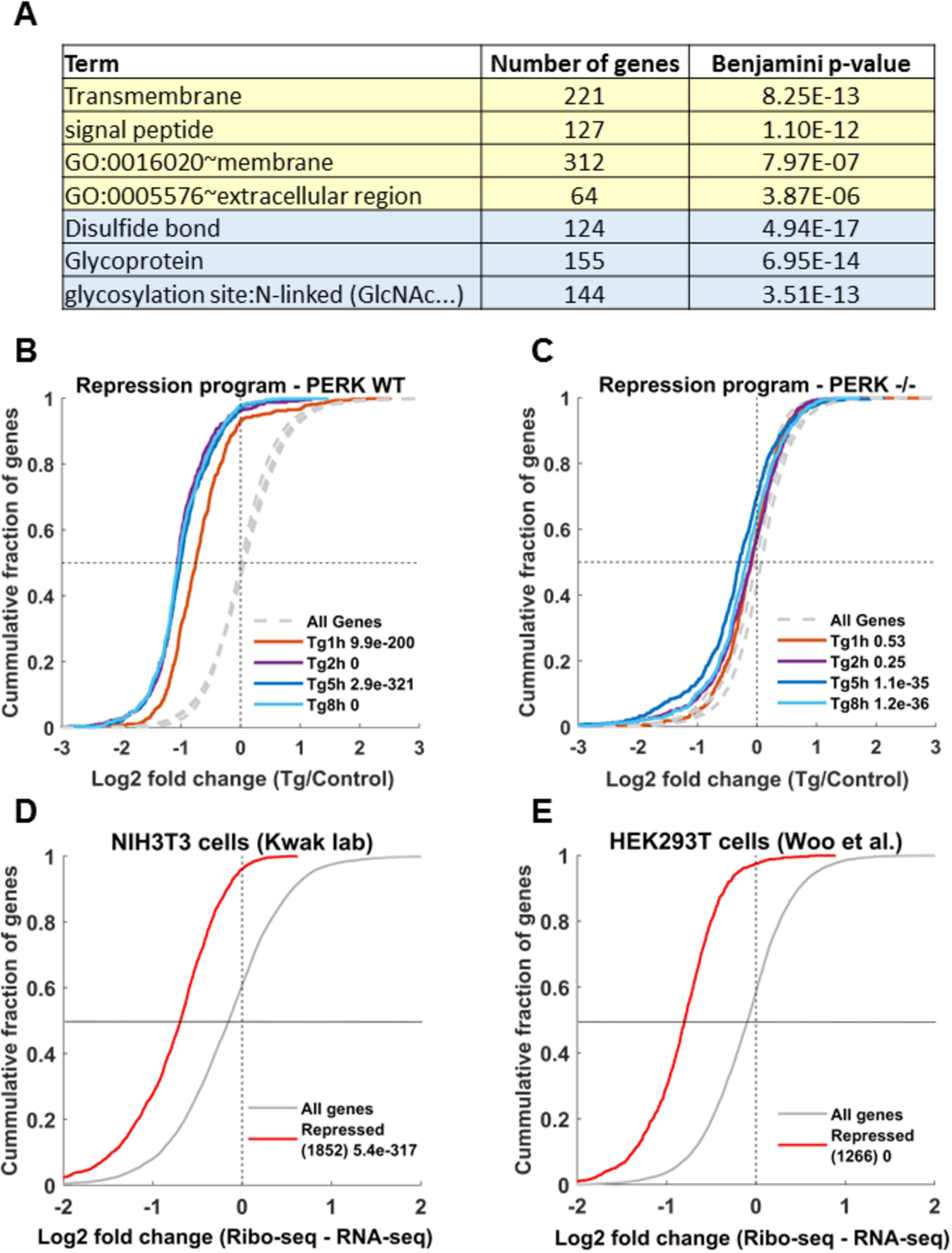
Widespread PERK-dependent repression of ER targets during ER stress. (A) Functional enrichment analysis of the ER stress repression gene expression program was performed using DAVID^38^. The enriched terms converged upon two main categories: membrane, transmembrane and signal peptide encoding proteins, and glycoproteins and disulfide-bond containing proteins, both describing sets of ER targets. (B,C) CDF plots demonstrate the cumulative fraction of repression program genes (y-axis), as a function of their log2 fold change (LFC) (x-axis) relative to control in (B) PERK WT and (C) PERK -/- cells. Dashed grey lines depict the background LFC distributions of all expressed genes in the different timepoint, t-test p-values between each distribution and its respective background are indicated. (D-E) ER stress-mediated repression shows a major translational regulatory component in both NIH3T3 cells (D) and HEK293T cells (E). CDF plots of the difference between the LFC at the translation level (Ribo-seq) and the LFC at the mRNA level (RNA-seq) are markedly shifted showing that repressed mRNAs are largely repressed at the level of translation. Repressed genes mRNAs defined as mRNAs with translation LFC of 1.5 fold repression or more. Both groups were highly enriched for ER targets (Table S2). This analysis was robust to various choices of repression parameters (Fig. S6J-M).

Next, we examined whether the repression of gene expression was dependent on PERK. In sharp contrast to the PERK WT cells (Fig. 5B), no downregulation was observed for the repression program genes in the PERK -/- cells at the early timepoints (Fig. 5C). At the later stage, a significant, yet minimal, repression was observed for these genes; with a median repression of 18% and 12% in the 5h and 8h timepoints respectively in PERK -/- cells, compared to a median of 50% and 51% repression in PERK WT cells (Fig. 4E,F).

These results indicate that, in addition to the global repression of protein synthesis caused by the PERK-induced eIF2α phosphorylation, ER targets are significantly and selectively further repressed, in a PERK dependent manner.

### Translational regulation plays a major role in the repression of ER targets during ER stress

As ribosome footprint profiling provides an integrated output of mRNA-level and translation-level changes, we further asked whether the PERK-dependent repression of ER targets that we identified occurs at the mRNA level or at the level of translation. Guan et al.^9^ have reported that ER-translated mRNAs were repressed in ER stress, with different subsets regulated at the mRNA level and at the level of translation. To further characterize the relationship between mRNA-level and translation-level regulation for the case of ER target repression, we separated genes that were repressed at 1h in the Guan et al. dataset into two groups: genes that showed repression both at the mRNA and the translational levels, and genes whose repression was mainly at the level of translation (Fig. S6A, see Materials and Methods). We note that, overall, the Guan et al. dataset showed a strong translational response at 1h, while at 16h of ER stress, mRNA levels and translation levels are highly correlated (Pearson R^2^ of 0.54, compared to 0.22 at 1h, Fig. S6A-C). Importantly, functional enrichment analysis of both groups showed enrichment for ER targets (Table S2A,B). Further examination of these groups at 16h of ER stress showed that their mode of regulation was largely maintained: while the first group was still repressed both at the mRNA and the translational levels, the second group remained mostly translationally repressed (Fig. S6B-E).

Next, we analyzed two additional datasets of mRNA-seq and ribosome footprint profiling (Ribo-seq), generated in NIH3T3 (Kwak lab, GSE103667) and HEK293T cells^18^ treated with a short ER stress, 1.5h and 2h respectively, using Tg. We first similarly defined groups of genes that were either repressed both at the mRNA level and translationally, or only translationally (see Materials and Methods). Importantly, in both datasets, a much smaller fraction of genes showed mRNA-level repression (15.3% and 6.6% of the repressed genes at a 1.5 fold cutoff, respectively), while the vast majority were translationally repressed (Fig. 5D,E, Fig. S6F-I, see Materials and Methods). Additionally, in both datasets, both translationally repressed genes and genes inhibited at the mRNA- and translation levels, were highly enriched for ER targets (Table S2C-F). We note that this analysis was robust to the choice of repression cutoff (see Materials and Methods, Fig. S6J-M, Table S2G-N).

These results further support the notion that the repression of ER targets upon ER stress is a widespread phenomenon, occurring at the level of translation for a substantial part of the genes, with mRNA-level contribution for a subset of the genes.

### PERK-mediated repression of ER targets during ER stress occurs in other cell types

As we have shown above, the repression of ER targets occurs at the level of translation, as well as at the mRNA level for a subset of genes. This fact allowed us to examine whether we can observe any downregulation of our repression gene expression program, which was highly enriched with ER targets, while examining other mRNA expression profiling datasets.

We first wanted to verify that the repression gene expression program we identified above was general to ER stress, and is not Thapsigargin specific. Analysis of two additional mRNA expression profiling datasets of MEFs treated with Tunicamycin (TM), for short, intermediate^19^ or long^12^ treatments, showed a mild yet highly significant downregulation of our identified repression program (Fig. S7A-C). Furthermore, analysis of mRNA expression profiling from DTT treated MEFs (Kaufman lab, GSE84450) also recapitulated this repression (Fig. S7D).

Therefore, our identified gene repression program was not Tg specific, but rather a general characteristic of the response to ER stress.

We then analyzed several additional mRNA expression profiling datasets to see if the repression program can be generalized to other specialized cell types, beyond mouse fibroblasts and human HEK293T cells (Fig. 5E, S6I,K,M). Indeed, we found that significant inhibition of the repression program was recapitulated in intestinal stem cells subject to 4h of TM^20^ (Fig. S7E), and in TM treated glioma-derived stem cells, and glioblastoma cells (Dorsey lab, GSE102505) (Fig. S7F,G), but not in livers treated with 6h of TM^21^ (Fig. S7H).

These analyses indicate that overall, the repression gene expression program is general, and occurs across many different, although not all, cell types.

### PERK-mediated repression of ER targets during ER stress largely involves eIF2α phosphorylation, and is partly dependent on ATF4

We then turned to explore the potential role of regulators downstream to PERK in the mediation of ER target repression. To that end, we first analyzed a dataset of mRNA expression from a long ER stress treatment (TM, 12h), performed in either WT, PERK -/- or eIF2α-S51A MEFs (Oyadomari lab, GSE49598). We observed a mild but significant repression at the level of mRNA in WT MEFs, both for our defined repression program (Fig. S8A), as well as for ER targets (signal peptide containing proteins, Fig. S8D). This repression was completely abrogated in PERK -/- MEFs (Fig. S8B,E), consistent with our ribosome footprint profiling data above. Furthermore, eIF2α-S51A MEFs also showed no repression of ER targets, as well as of the repression program (Fig. S8C,F).

Interestingly, our analysis of a dataset of Arginine deprivation in HCT116 and HEK293T cells^22^, a condition that is known to lead to phosphorylation of eIF2α through GCN2, also found a mild yet significant downregulation of the repression program (Fig. S8G-I). However, this repression did not depend on GCN2 (Fig. S8J), since it remained slight but significant in GCN2 knockout HEK293T cells.

These results indicate that eIF2α phosphorylation downstream of PERK plays a major role in the preferential repression of ER targets (see discussion below).

The ATF4 transcription factor is an important effector in the ER stress response downstream to PERK, in an eIF2α phosphorylation dependent manner (Fig. S1B). Since we observed the repression of ER targets as early as 1h of stress, and a large part of it was translational, it is less likely to be dependent on ATF4. However, since mRNA-level repression of some ER targets does occur (Fig. S6), we sought to further examine whether ATF4 plays a role in this repression. Analysis of a dataset of 8h ER stress (TM treatment) in either WT or ATF4-/- MEFs^12^ showed a significant downregulation of our identified repression program in both WT and ATF4-/- cells (Fig. S8K,L). These results indicate that the PERK-mediated repression program is largely independent on ATF4. Nevertheless, when we examined the set of ER targets (signal peptide encoding mRNAs), we observed a partial relief in the repression of a subset of them in ATF4-/- MEFs (Fig. S8M,N). Thus, the repression of a subset of ER targets is partially dependent on ATF4.

### Additional pathways subject to PERK-mediated repression

To ask whether the repression cluster contains additional pathways, we removed all 265 ER targets from this cluster, leaving a gene set of 393 repressed genes, and repeated the functional enrichment analysis. We found that this repressed set is enriched for cyclins (Table S1D). This enrichment in repressed cyclins fits well with the cell cycle arrest that occurs upon ER-stress^23^. Further analysis showed that cyclins were repressed as a group (Fig. S9A), and this repression did not occur in PERK -/- cells. Additionally, we observed an enrichment for LSM domain proteins (Table S1D), a family of RNA-binding proteins. For this group as well, repression was PERK dependent (Fig. S9B).

### PERK attenuates a subset of XBP1 and ATF6 targets, revealing a complex interplay between the UPR arms

At the late stages, transcriptional responses mediated by the ATF6 and IRE1-XBP1 arms of the UPR are expected to take effect. We found that the expression levels of both XBP1 and ATF6 were induced by ER stress, and this induction was partly diminished (40% less induction) in PERK -/- cells (Fig. 1E, and S10), in agreement with previous reports^21^. We wanted to better understand the potential interplay between the XBP1- and ATF6- mediated response pathways and PERK. We therefore examined the expression pattern of bona-fide XBP1 and ATF6 target gene sets, which were originally defined by Shoulders et al.^24^ without eliciting ER stress, within our data. In the early timepoints, the expression of XBP1 and ATF6 targets was largely unchanged (Fig. S11A-C). This is consistent with the delayed accumulation in their transcriptional output. In the late timepoints, however, we found that while 74% of XBP1-ATF6 targets were induced in PERK -/- cells following ER stress, only 56% were induced in PERK WT cells (at 8h, Fig. 6A,B). Similar trends were observed when we considered XBP1 targets and ATF6 targets separately (Fig. S11D,E). Thus, it seems that PERK might have an inhibitory effect on a subset of XBP1 and ATF6 target genes.

**Figure 6.**
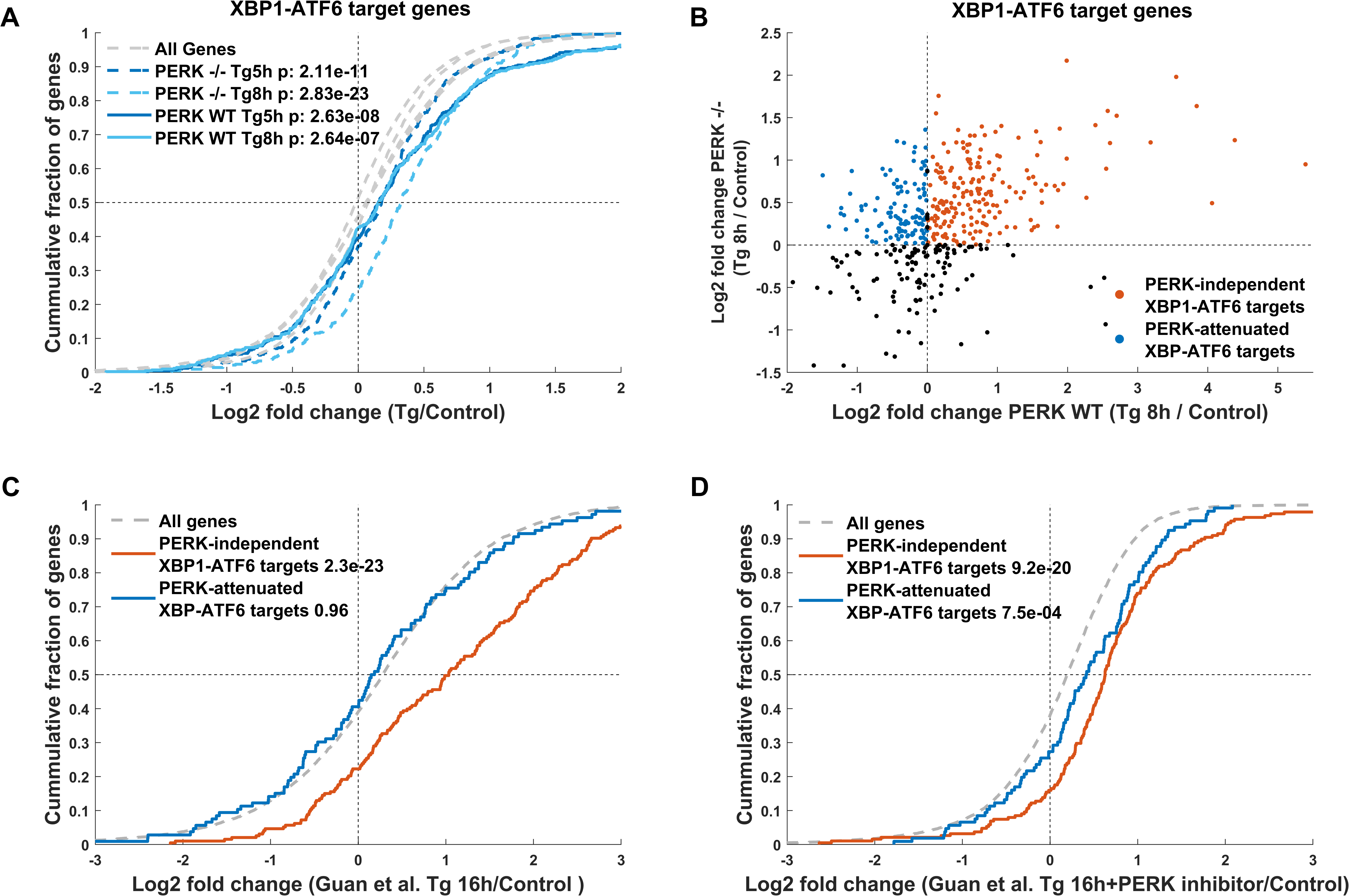
PERK-mediated attenuation of a subset of XBP1-ATF6 targets during the late stages of ER stress. XBP1-ATF6 target genes, as defined by Shoulders et al.^24^, were examined for their expression during ER stress. (A) CDF plot of the log2 fold changes (LFCs) of this gene set in the late timepoints demonstrates that these genes are largely induced in PERK -/- cells (dashed blue lines), while a smaller subset of genes is induced in PERK WT cells (solid blue lines). Grey dashed lines indicate the background distributions of the LFCs of all expressed genes in the different conditions, t-test p-values between each distribution and its respective background are indicated. (B) Scatterplot of the 8h Tg LFCs in PERK WT (x-axis) vs. PERK -/- (y-axis) demonstrate that the majority of XBP1-ATF6 targets are induced in PERK -/- cells, and defines two subsets of XBP1-ATF6 targets: targets that are induced in both PERK WT and -/- MEFs, PERK-independent XBP1-ATF6 targets (red circles), and PERK-attenuated XBP1-ATF6 targets (blue circles); induced in PERK -/- but not in PERK WT cells. (C,D) CDF plots show that PERK-independent XBP1-ATF6 targets are induced in the Guan et al. polysome-associated mRNA profiling dataset^9^ after 16h of Tg treatment (C, red curve) irrespective of PERK inhibition (D, red curve), while PERK-attenuated XBP1-ATF6 targets were unchanged at 16h Tg treatment (C, blue curve), and induced upon PERK inhibition (D, blue curve).

To better understand the effect of PERK on XBP1-ATF6 targets, we further defined two gene sets: PERK independent XBP1-ATF6 targets, i.e. XBP1-ATF6 targets that were induced in both PERK WT and PERK -/- cells (196 genes, Fig. 6B, red circles), and PERK-attenuated XBP1-ATF6 targets, targets that were induced only in PERK -/- cells but not in PERK WT cells (109 genes, Fig. 6B, blue circles).

We further examined the expression of these two gene sets in the Guan et al. polysome-seq data^9^, where MEFs have been ER stressed for 16h, with or without the presence of a PERK inhibitor for the last 4h of the stress. Our analysis revealed that, consistent with our data above, PERK-independent XBP1-ATF6 targets were induced at 16h of ER stress irrespective of PERK inhibition (Fig. 6C,D, red curve). The PERK-attenuated XBP1- ATF6 targets, however, showed no induction at 16h post stress (Fig. 6C, blue curve). Nevertheless, the expression pattern of these genes in the PERK inhibitor treated cells demonstrated that PERK inhibition alleviated this attenuation (Fig. 6D, blue curve), further substantiating that this subset of XBP1-ATF6 targets is subject to PERK-mediated attenuation.

Similar trends were observed when we examined XBP1 targets separately (Fig. S11F-H).

To understand if this PERK-mediated attenuation occurs at the level of transcription or translation, we analyzed mRNA expression data from TM treated WT or PERK -/- MEFs (12h, Oyadomari lab, GSE49598). Here too, we observed the same relief in PERK-mediated inhibition of the subset of PERK-attenuated genes we identified above (Fig. S12A-B), indicating that transcriptional regulation is involved in this attenuating effect. Moreover, our analyses found that similar trends of induction vs. attenuation for the PERK-independent and PERK-attenuated XBP1-ATF6 targets defined above, were evident and highly significant in many mRNA expression datasets, including response to TM^19^ (Fig. S12D) and DTT (Kaufman lab, GSE84450, Fig. S12E), as well as in other cell types, including Tg treated intestinal stem cells^20^ (Fig. S12F), and human derived glial cells (24h of TM treatment, Dorsey lab, GSE102505, Fig. S12G).

Nevertheless, examination of mRNA expression data from Guan et al. MEFs^9^, where PERK inhibitor was added only during the last 4h of a 16h Tg treatment, showed a more complex picture. While these cells showed alleviation of attenuation at the translation level (Fig. 6D), no relief was observed at the mRNA level (Fig. S12H,I).

Taken together, our analyses suggest that the PERK-mediated attenuation of a subset of XBP1-ATF6 targets contains both translational and transcriptional components, as initial relief in attenuation is observed at 4h of PERK inhibitor only at the level of translation. At longer durations of ER stress, transcriptional regulation downstream of PERK seems to play a major role in the attenuation of this subset of XBP1-ATF6 targets.

Analysis of the potential contribution of PERK pathway effectors showed that eIF2α- S51A MEFs (Oyadomari lab, GSE49598) largely recapitulate the relief of the PERK-mediated attenuation of this subset of XBP1-ATF6 targets (Fig. S12C) observed in PERK -/- MEFs. Furthermore, in ATF4 -/- MEFs^12^ this attenuation also showed significant alleviation (Fig. S12J,K). These results indicate that eIF2α phosphorylation is largely involved in mediating the PERK-dependent attenuation of this subset of XBP1-ATF6 targets, with a significant contribution of the ATF4 branch to the phenomenon, in line with the combined transcriptional-translational regulatory effect downstream of PERK we found above.

The fact that this phenomenon had similar characteristics to the PERK-mediated repression of ER targets, prompted us to speculate that these two phenomena might be related. Indeed, functional enrichment analysis showed that many of these PERK-attenuated XBP1-ATF6 targets are ER targets (Table S3B,C). Therefore, it is possible that this subset of targets is subject to the PERK-dependent ER targets repression program described above. The PERK-attenuated XBP1-ATF6 target set was selectively enriched with sterol and lipid biosynthesis genes (Table S3B), which are ER residents, and their repression following ER stress has been previously reported^25^. ER targets were also enriched within the PERK-independent set (Table S3A,C). There, however, selectively enriched ER targets included ER stress response pathway genes (Table S3A), indicating that classical ER stress response regulators are not subject to attenuating regulation via PERK.

Together, our analyses revealed a major gene expression program including PERK-mediated repression of ER targets. This program also attenuates a subset of XBP1-ATF6 targets, unravelling another level of the complex interplay between the different arms of the ER stress response.

## Discussion

The ribosome profiling high-throughput data presented in our study provides a genome-wide temporal view of ER stress gene expression programs, integrating transcription and translational outputs, in both WT and PERK -/- MEFs. Our data and analyses revealed composite dynamics of the integrated expression response to ER stress, orchestrated by PERK, and showed a complex combination of gene repression and induction under ER-stress.

Our data suggests that the impairment in specific gene activation, as well as repression, in the absence of PERK are very widespread, and unravel a complex interplay between the three arms of the ER stress response.

Wang et al.^26^ has reported a strong and immediate global translational repression within ten minutes after pharmacological ER-stress induction, using nascent chain translation live imaging. The polysome profiling above (Fig. 1A,B) demonstrated that this repression was markedly noticeable 1h post Tg treatment and was gradually relieved.

Our findings capture a strong, selectively enhanced, repressive gene expression program following ER stress and demonstrate its impact on ER targeted proteins (Fig. 5), which seems to be an additional hallmark of the ER stress response. More specifically, we found that proteins undergoing ER-dependent post-translational modifications such as glycosylation and disulfide-bonds, as well as transmembrane and membranal proteins are enriched among genes repressed under ER-stress. Importantly, we have shown that this repression occurs largely at the level of translation, with a small contribution of mRNA-level repression (Fig. 5D,E, S6).

Several microRNAs have been shown to be induced in response to ER stress^27,28^. However, examination of the putative targets of these miRNAs did not explain the repression of ER targets observed (data not shown). Rapid changes in translation-dependent compartmentalization of ER-target protein synthesis has been previously demonstrated, with an increase in the proportional translation of ER targets by cytosolic ribosomes^10^. This change in translation compartmentalization was shown to be transient, with a robust change at 30 minutes post Tg treatment, and an almost complete reversal by 1h following ER-stress induction^10^. Significant repression in ER stress has been later reported for these genes^9^. Here we show that there is major translational repression of ER targets, which is highly PERK dependent. Importantly, even though it has an mRNA-level component, translation is the major regulatory level for this phenomenon, and thus it was strongly evident from our ribosome footprint profiling data analysis. It is possible that the re-localization of ER targets^10^ may be an initiating step in the cellular response to ER stress, with a second, yet to be discovered, PERK-mediated mechanism of repression. Moreover, our analyses point to eIF2α phosphorylation as critical for this repression. One possible model that would be consistent with the data is that PERK-mediated phosphorylation of eIF2α subunits primarily affects the translational machinery pool at the vicinity of the ER, thus affecting mainly ER targets. Our analysis of ribosome footprint profiling data from Arginine deprived cells^22^ has shown that while this stress elicits mild ER target repression, this repression was GCN2-independent. It is therefore possible that in the case of GCN2, eIF2α phosphorylation, which occurs throughout the cytoplasm, does not lead to a GCN2-dependent repression of ER targets. However, whether indeed this model underlies PERK-mediated repression of ER targets remains to be explored.

Past research has indicated Cyclin D1 to be translationally repressed during ER-stress in a PERK dependent manner ^23^. Interestingly, we observed a wider PERK-mediated repression of cyclins expression, including cyclins from the D,E,F,G and I families, thereby expanding existing knowledge regarding PERK mediated cell-cycle arrest. The cell-cycle arrest has immense implications, which may be essential for cell survival under chronic stress^23^. Furthermore, we found a PERK-dependent LSM domain protein repression, pointing towards a possible regulation at the level of RNA processing. LSM-proteins are known to aggregate within processing-bodies under various stress conditions altering mRNA homeostasis^29^. In addition, we see an early transient PERK dependent induction of DNA-binding proteins, mainly zinc-finger and histones, in agreement with previous reports^10^. These yet to be explored changes may be related to ER stress-mediated chromatin changes. For example, ATF4 target gene activation has been linked to the histone lysine demethylase KDM4C epigenetic rewiring in cancer^30^.

Previous studies have underlined PERK reinforcement of both the IRE1-XBP1 and ATF6 UPR arms^21,31^. It has been shown previously that XBP1-spliced accumulation is PERK dependent^31^. Furthermore, the transcription of selected ATF6 targets is augmented 6h after ER-stress initiation in the liver, and the processing of ATF6 has been shown to be dependent on ATF4^21^. Recent research has reported PERK-mediated IRE1 inhibition via the RPAP2 phosphatase, which has a role in diverting adaptive ER-stress response towards apoptosis after 16h of stress^32^. Consistently, here we observed that both XBP1 and ATF6 expression are induced in a temporal manner following ER stress, and their induction is partially reduced in the absence of PERK (Fig. 1E, Fig. S10). Thus, several reinforcing positive interactions between PERK and other UPR arms have been demonstrated. Nevertheless, we report here that PERK mediates an attenuating effect on a subset of XBP1 and ATF6 target genes (Fig. 6, S11). As many of these genes are ER targets (Table S3), they could be subject to the same mechanism of PERK-mediated repression of ER targets we highlight above. Notably, sterol biosynthetic genes, which are directly upregulated by XBP1-ATF6 activation, were specifically attenuated in a PERK-dependent manner (Table S3B). Indeed, previous findings have linked the UPR to down-regulation of the sterol biogenesis pathway, and subsequently cholesterol levels, via suppression of SREBP^25^. This PERK-mediated attenuation that we found was not gradually relieved at 8h (Fig. 6A), nor by 16h of ER stress, as seen from our re-analysis of the dataset by Guan et al.^9^ (Fig. 6C). The repression gene expression program shows a similar trend (Fig. 5B, S3C). Furthermore, both phenomena show similar characteristics of eIF2α phosphorylation dependence and a smaller yet significant contribution via ATF4. This raises the possibility that these two phenomena might indeed be related, such that ER target repression explains the PERK-mediated attenuation of the subset of XBP1- ATF6 targets. While repression of ER targets is primarily translational and starts early, and transcriptional effects kick in later on, and then this subset of the transcriptional UPR arms is attenuated. While future studies will unravel the purpose of this attenuating affect, and its role in the adaptation to ER stress, it is clear that the complex cross-talk between the three arms of the UPR, the interplay between transcriptional and translational regulation, and the function of PERK in this interplay, play a critical role in shaping the cellular gene expression program in response to ER stress.

## Materials and Methods

### Cell culture and stress

Mouse fibroblast NIH3T3 cells were grown in DMEM with 10% FBS and pen-strep. MEFs were grown in DMEM with 10% FBS supplemented with 0.1 mM Non-Essential Amino Acids (NEAA) and 0.05mM beta-Mercaptoethanol. NIH3T3 cells or MEFs were plated at low density (2 million cells in 15 cm plate one day prior to harvesting), and treated with Thapsigargin (at 1µM or 200nM for MEFs and NIH3T3 cells respectively) for short (1h or 2h) or long (5h and 8h for MEFs, 7h for NIH3T3) durations, to induce early or late ER stress responses.

### Ribosome footprint profiling and Polysome profiling

Ribosome footprint profiling was performed as in Shalgi et al.^16^, with the following modification: cells were grown and treated with Tg for the indicated times, then trypsinized, pelleted by 5 minutes at 1000RPM centrifugation at 4°c, and flash frozen in LN2, without prior Cycloheximide treatment. Cycloheximide was also omitted from the buffers. Polysome profiling was performed as in Shalgi et al.^16^.

### Data analysis

Ribosome footprint profiling provided quantification of number of ribosomes per transcript in each timepoint, resulting in the quantification of protein synthesis levels. We note that the protein synthesis levels are relative, rather than absolute, since the overall number of translating ribosomes is decreased in response to ER stress in WT MEFs, due to eIF2α phosphorylation^8^, as also observed in Fig. 1. Nevertheless, we were interested in characterizing, given the limited pool of overall translating ribosomes, which genes and which pathways are relatively more enhanced or further repressed under each condition.

Ribosome footprint profiling data was mapped to mm10 version of the genome using RefSeq CDSs. Transcripts shorter that 100 nucleotides were filtered out and 30 nucleotides were clipped form the start and end of each CDS, similarly to Ingolia et al.^33^. Ribosome footprint sequences were trimmed to maximal size of 34 and polyA sequences from the polyA tailing step were removes, such that footprints of lengths 22-34 were considered. FASTQ files were then filtered for rRNAs and tRNAs using STAR^34^. Expression levels were quantified using RSEM^35^ after mapping to clipped CDSs using Bowtie2^36^ to produce transcript and gene level TPM (Transcript Per Million) values (see Table S5).

Lowly expressed genes with an average TPM below 2 across all samples were filtered out and expression of all other genes was thresholded to a TPM of 4. For each experimental group (PERK WT or -/- MEFs, or NIH3T3 cells), fold changes were calculated as a ratio between each condition and its respective control. Regulated genes were designated as genes which changed at least two-fold relative to their designated controls.

PCA was generated using R DESEQ2^37^, using the function plotPCA. Hierarchical clustering of genes was done based on spearman correlation using the clustergram MATLAB function.

Additional RNA-seq ER-stress datasets (detailed in Table S4) were downloaded from the GEO database. RNA-Seq reads were filtered for rRNAs using STAR^34^. The remaining reads were mapped to hg19 or mm10 versions of the human and mouse genomes, respectively, using STAR^34^. Expression levels were quantified using RSEM^35^. TPM values were averaged between sample replicates.

Ribosome footprint profiling data were processed as described above (mapped to hg19 or mm10 versions of the human and mouse genomes, respectively).

Microarray ER stress datasets (detailed in Table S4) were downloaded from GEO. Fold changes for the dataset GSE54581 were derived using the Affymetrix transcriptome analysis console, with CEL files as input. Fold changes for the datasets GSE49598 and GSE84989 were generated using normalized gene expression signals as provided by GEO. Replicates from each condition of interest were averaged, and gene expression fold changes were calculated.

To characterize the relationship between mRNA-level and translation-level regulation for the case of ER target repression, we separated the repressed genes (1.5 fold) in three datasets (Guan et al.^9^, Woo et al.^18^, and Kwak lab NIH3T3 cells) into two groups: genes that showed repression both at the mRNA and the translational level (mRNA-level repression and translation-level repression 1.5 fold or more), and genes whose repression was mainly at the level of translation (mRNA-level repression less than 1.5 fold and translation-level repression more than 1.5 fold). We examined the number of genes in each group as well as the difference between log2 fold change in translation-level and log2 fold change in mRNA-level. We repeated the above analyses using a range of threshold values from 1.3-2.1 and verified that the results were indeed robust to the choice of threshold (Fig. S6J-M). Additionally, we repeated the functional enrichment analysis using sets with different thresholds for these two datasets, and all translationally repressed gene groups, as well as mRNA and translationally repressed gene groups, were found to be significantly enriched for ER targets (Table S2).

CDF plots were used to compare the cumulative distributions of log2 fold changes (LFCs) in the expression of gene subsets compared to their respective control, to the overall distribution of LFCs of the entire transcriptome (expressed genes, defined as above). Significance for different gene groups was calculated using the student t-test statistical test.

Pathway enrichment analyses were conducted using DAVID 6.8 functional annotation tool ^38^, using all expressed genes as background. Pathways with Benjamini corrected p-value <0.05 were designated as enriched.

Signal peptide encoding mRNAs were defined using the signalP program^39^.

XBP1-ATF6 targets were defined by Shoulders et al.^24^ following specific activation of XBP1, ATF6, or both of them together (termed XBP1-ATF6 targets). We further filtered these sets for genes that were not expressed in our cells (TPM<2). PERK-independent XBP1-ATF6 targets were defined as XBP1-ATF6 targets that were upregulated in both PERK WT and PERK -/- following 8 hours of Tg, and PERK-attenuated XBP1-ATF6 targets were defined as XBP1-ATF6 targets up-regulated in PERK -/- and down-regulated in PERK WT. These sets were further subject to functional enrichment analysis using DAVID, and the resulting enriched terms (Benjamini corrected p<0.05) were examined for the uniqueness in only one of the sets (Tables S3A,B) or intersected in both sets (Table S3C).

## Supporting information

## Acknowledgements

We would like to thank Ruth Hershberg for useful comments and critical reading of the manuscript, Anatoly Meller for critical reading of the manuscript, and Mals Mariappan for useful discussion of the results. This project has received funding from the European Research Council under the European Union’s Horizon 2020 programme Grant 677776.

## Author contributions

RS conceived and supervised the study. RS and CBB designed the study. RS performed the experiments, NG designed and performed all data analysis with the help of NS.

## Data availability

Original and processed data were deposited in GEO under GSE118660. Processed data is also available in Table S5. Additional datasets used are listed in Table S4.

## Electronic supplementary material

Supplementary Figures S1-S12

Supplementary Tables S1

Supplementary Tables S2

Supplementary Tables S3

Supplementary Tables S4

Supplementary Tables S5

## References

1 Reid, D. W. & Nicchitta, C. V. Diversity and selectivity in mRNA translation on the endoplasmic reticulum. Nature reviews. Molecular cell biology 16, 221–231, doi:10.1038/nrm3958 (2015).

2 Ron, D. & Walter, P. Signal integration in the endoplasmic reticulum unfolded protein response. Nature reviews. Molecular cell biology 8, 519–529, doi:10.1038/nrm2199 (2007).

3 Feige, M. J. & Hendershot, L. M. Disulfide bonds in ER protein folding and homeostasis. Current Opinion in Cell Biology 23, 167–175, doi:https://doi.org/10.1016/j.ceb.2010.10.012 (2011).

4 Sonenberg, N. & Hinnebusch, A. G. Regulation of Translation Initiation in Eukaryotes: Mechanisms and Biological Targets. Cell 136, 731–745, doi:10.1016/j.cell.2009.01.042 (2009).

5 Hetz, C., Chevet, E. & Oakes, S. A. Proteostasis control by the unfolded protein response. Nature cell biology 17, 829–838, doi:10.1038/ncb3184 (2015).

6 Hollien, J. Evolution of the unfolded protein response. Biochimica et biophysica acta 1833, 2458–2463, doi:10.1016/j.bbamcr.2013.01.016 (2013).

7 Pavitt, G. D. & Ron, D. New insights into translational regulation in the endoplasmic reticulum unfolded protein response. Cold Spring Harbor perspectives in biology 4, doi:10.1101/cshperspect.a012278 (2012).

8 Sonenberg, N. & Hinnebusch, A. G. Regulation of translation initiation in eukaryotes: mechanisms and biological targets. Cell 136, 731–745, doi:S0092- 8674(09)00090-7 10.1016/j.cell.2009.01.042 (2009).

9 Guan, B. J. et al. A Unique ISR Program Determines Cellular Responses to Chronic Stress. Mol Cell 68, 885–900 e886, doi:10.1016/j.molcel.2017.11.007 (2017).

10 Reid, D. W., Chen, Q., Tay, A. S., Shenolikar, S. & Nicchitta, C. V. The unfolded protein response triggers selective mRNA release from the endoplasmic reticulum. Cell 158, 1362–1374, doi:10.1016/j.cell.2014.08.012 (2014).

11 Harding, H. P., Zhang, Y., Bertolotti, A., Zeng, H. & Ron, D. Perk Is Essential for Translational Regulation and Cell Survival during the Unfolded Protein Response. Molecular cell 5, 897–904, doi:https://doi.org/10.1016/S1097-2765(00)80330-5 (2000).

12 Han, J. et al. ER-stress-induced transcriptional regulation increases protein synthesis leading to cell death. Nat Cell Biol 15, 481–490, doi:10.1038/ncb2738 (2013).

13 Harding, H. P. et al. An integrated stress response regulates amino acid metabolism and resistance to oxidative stress. Molecular cell 11, 619–633 (2003).

14 Ingolia, N. T., Ghaemmaghami, S., Newman, J. R. & Weissman, J. S. Genome-wide analysis in vivo of translation with nucleotide resolution using ribosome profiling. science 324, 218–223 (2009).

15 Vattem, K.M. & Wek, R. C. Reinitiation involving upstream ORFs regulates ATF4 mRNA translation in mammalian cells. Proc Natl Acad Sci U S A 101, 11269–11274, doi:10.1073/pnas.0400541101 (2004).

16 Shalgi, R. et al. Widespread Regulation of Translation by Elongation Pausing in Heat Shock. Molecular cell 49, 439–452, doi:10.1016/j.molcel.2012.11.028 (2013).

17 Liu, B., Han, Y. & Qian, S.-B. Cotranslational Response to Proteotoxic Stress by Elongation Pausing of Ribosomes. Molecular cell 49, 453–463, doi:10.1016/j.molcel.2012.12.001 (2013).

18 Woo, Y. M. et al. TED-Seq Identifies the Dynamics of Poly(A) Length during ER Stress. Cell Rep 24, 3630–3641 e3637, doi:10.1016/j.celrep.2018.08.084 (2018).

19 Miyazaki, Y., Chen, L. C., Chu, B. W., Swigut, T. & Wandless, T. J. Distinct transcriptional responses elicited by unfolded nuclear or cytoplasmic protein in mammalian cells. Elife 4, doi:10.7554/eLife.07687 (2015).

20 Tsalikis, J. et al. The transcriptional and splicing landscape of intestinal organoids undergoing nutrient starvation or endoplasmic reticulum stress. BMC Genomics 17, 680, doi:10.1186/s12864-016-2999-1 (2016).

21 Teske, B. F. et al. The eIF2 kinase PERK and the integrated stress response facilitate activation of ATF6 during endoplasmic reticulum stress. Mol Biol Cell 22, 4390–4405, doi:10.1091/mbc.E11-06-0510 (2011).

22 Darnell, A. M., Subramaniam, A. R. & O’Shea, E. K. Translational Control through Differential Ribosome Pausing during Amino Acid Limitation in Mammalian Cells. Mol Cell 71, 229–243 e211, doi:10.1016/j.molcel.2018.06.041 (2018).

23 Brewer, J. W. & Diehl, J. A. PERK mediates cell-cycle exit during the mammalian unfolded protein response. Proc Natl Acad Sci U S A 97, 12625–12630, doi:10.1073/pnas.220247197 (2000).

24 Shoulders, M. D. et al. Stress-independent activation of XBP1s and/or ATF6 reveals three functionally diverse ER proteostasis environments. Cell Rep 3, 1279–1292, doi:10.1016/j.celrep.2013.03.024 (2013).

25 Harding, H. P. et al. Bioactive small molecules reveal antagonism between the integrated stress response and sterol-regulated gene expression. Cell Metabolism 2, 361–371, doi:https://doi.org/10.1016/j.cmet.2005.11.005 (2005).

26 Wang, C., Han, B., Zhou, R. & Zhuang, X. Real-Time Imaging of Translation on Single mRNA Transcripts in Live Cells. Cell 165, 990–1001, doi:10.1016/j.cell.2016.04.040 (2016).

27 Chitnis, N. S. et al. miR-211 is a prosurvival microRNA that regulates chop expression in a PERK-dependent manner. Molecular cell 48, 353–364, doi:10.1016/j.molcel.2012.08.025 (2012).

28 Behrman, S., Acosta-Alvear, D. & Walter, P. A CHOP-regulated microRNA controls rhodopsin expression. The Journal of cell biology 192, 919–927, doi:10.1083/jcb.201010055 (2011).

29 Cornes, E. et al. Cytoplasmic LSM-1 protein regulates stress responses through the insulin/IGF-1 signaling pathway in Caenorhabditis elegans. Rna 21, 1544–1553, doi:10.1261/rna.052324.115 (2015).

30 Zhao, E. et al. KDM4C and ATF4 Cooperate in Transcriptional Control of Amino Acid Metabolism. Cell Rep 14, 506–519, doi:10.1016/j.celrep.2015.12.053 (2016).

31 Majumder, M. et al. A novel feedback loop regulates the response to endoplasmic reticulum stress via the cooperation of cytoplasmic splicing and mRNA translation. Molecular and cellular biology 32, 992–1003, doi:10.1128/MCB.06665-11 (2012).

32 Chang, T.-K. et al. Coordination between Two Branches of the Unfolded Protein Response Determines Apoptotic Cell Fate. Molecular cell 71, 629–636.e625, doi:10.1016/j.molcel.2018.06.038 (2018).

33 Ingolia, N. T., Brar, G. A., Rouskin, S., McGeachy, A. M. & Weissman, J. S. The ribosome profiling strategy for monitoring translation in vivo by deep sequencing of ribosome-protected mRNA fragments. Nature protocols 7, 1534–1550, doi:10.1038/nprot.2012.086 (2012).

34 Dobin, A. et al. STAR: ultrafast universal RNA-seq aligner. Bioinformatics 29, 15–21, doi:10.1093/bioinformatics/bts635 (2013).

35 Li, B. & Dewey, C. N. RSEM: accurate transcript quantification from RNA-Seq data with or without a reference genome. BMC Bioinformatics 12, 323, doi:10.1186/1471-2105-12-323 (2011).

36 Langmead, B. & Salzberg, S.L. Fast gapped-read alignment with Bowtie 2. Nature methods 9, 357–359, doi:10.1038/nmeth.1923 (2012).

37 Love, M. I., Huber, W. & Anders, S. Moderated estimation of fold change and dispersion for RNA-seq data with DESeq2. Genome Biol 15, 550, doi:10.1186/s13059-014-0550-8 (2014).

38 Jiao, X. et al. DAVID-WS: a stateful web service to facilitate gene/protein list analysis. Bioinformatics 28, 1805–1806, doi:10.1093/bioinformatics/bts251 (2012).

39 Nielsen, H. Predicting Secretory Proteins with SignalP. Methods Mol Biol 1611, 59–73, doi:10.1007/978-1-4939-7015-5_6 (2017).

