## Supplementary material for "Widespread PERK-dependent repression of ER targets in response to ER stress"

A

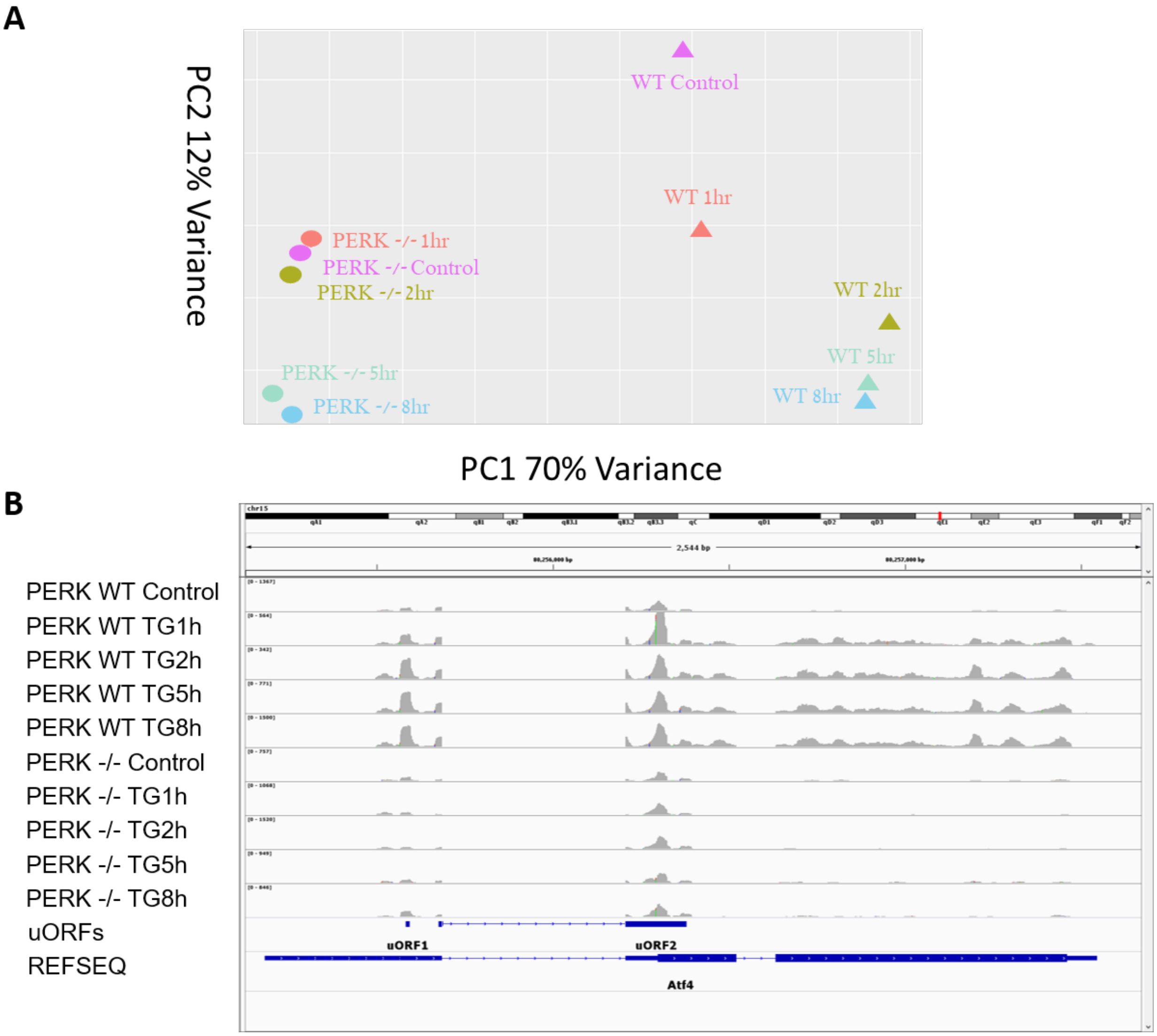


**Figure S1 – PCA analysis and ATF4 read density plots.**
(A) Principle Component Analysis shows a separation between PERK WT and -/- samples (PC1) and the temporal ER stress treatment (PC2). (B) ATF4 read density plot (in grey, using IGV) depicts uORFs and a PERK-dependent main ORF expression during ER stress. uORFs and main ORF transcripts are shown in blue below. Y-axis scales were normalized according to library depth to better illustrate the relative differences in the different conditions.

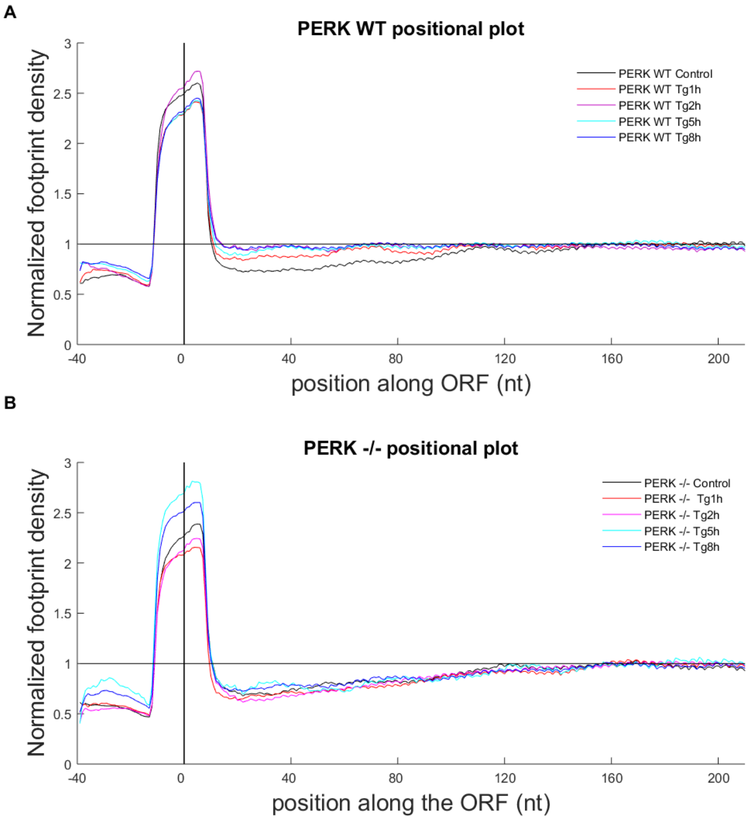


**Figure S2 - Positional plots of PERK WT and PERK -/- ribosome footprint profiling data**
Averaged normalized read density is plotted as a function of position for all expressed ORFs, produced similarly to Shalgi et al.[1] , 0 point indicated the AUG position.

Using KS test we observed a significant accumulation of ribosomes in the 5' ends of ORFs starting downstream to the AUG up until position 141 bases into the ORF ($p < 10^{-300}$ for all PERK WT ER stress samples compared to control, position and p-value determined by a KS-test). The magnitude of the accumulation was on average 25-35% in the different timpepoints in PERK WT MEFs (A). Notably, PERK -/- cells (B) did not show any ribosome accumulation.

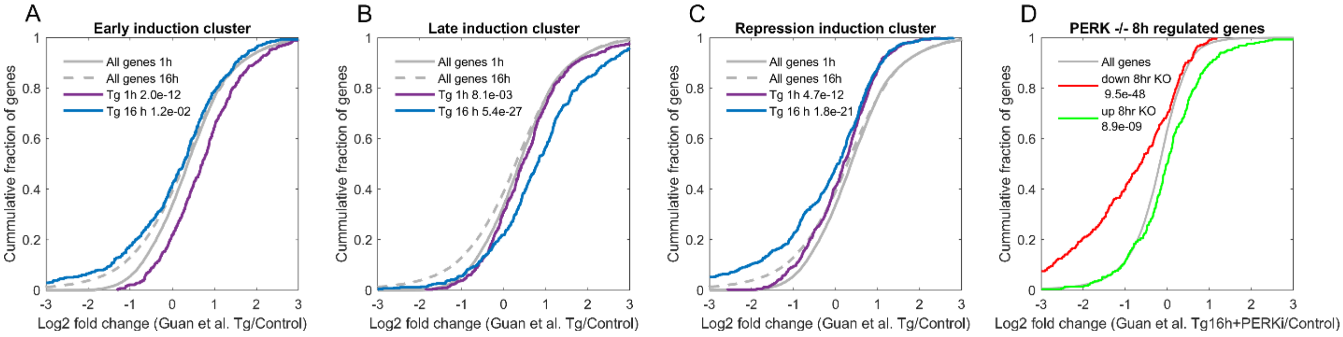


**Figure S3 - Major ER stress gene expression programs are recapitulated in a publicly available polysome-sequencing ER stress dataset.**

(A-C) Log2 fold changes (LFCs) of the genes from the three major ER stress gene expression programs, namely early induction (A), late induction (B) and repression (C), were calculated using a polysome-sequencing dataset from Guan et al.[2], of MEF cells treated with Tg for either 1h (purple) or 16h (blue). The analysis demonstrated that the three ER stress gene expression programs were recapitulated in this dataset. Grey solid and dashed lines indicate the background distributions of the LFCs of all expressed genes in the different timepoints, t-test p-values between each distribution and its respective background are indicated. (D) Induced (green) and repressed (red) genes were defined using the data of PERK -/- cells after 8h of Tg treatment from the current study (2 fold change compared to PERK -/- control cells). Data was extracted from Guan et al.[2] polysome-sequencing experiment done in MEFs following 16h of Tg treatment with a PERK inhibitor. The figure shows the CDF plot of the log2 fold change (LFC) of the two sets of induced and repressed PERK -/- Tg 8h genes, in the Guan et al. data of 16h ER stress with PERK inhibitor relative to untreated cells, and demonstrates that the induced set is also significantly induced in the Guan et al. polysome-seq data, while the repressed set is significantly repressed in the Guan et al. polysome-seq data. Grey line indicates the background distributions of the LFCs of all expressed genes, t-test p-values between the distributions and the background are indicated.

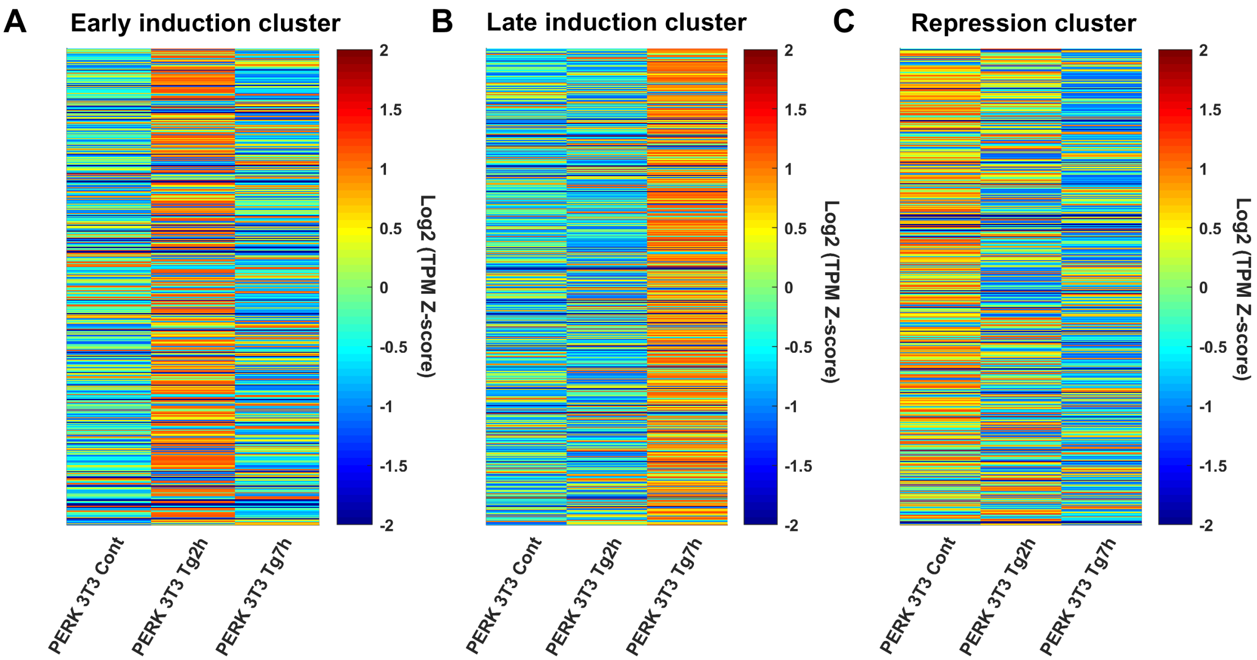


**Figure S4 - Heatmaps of PERK-dependent ER stress gene expression programs within NIH3T3 cells.**

Heatmaps of the Early induction **(A)**, Late induction **(B)** and Repression **(C)** gene expression programs were generated using TPMs calculated from NIH3T3 cells subject to either 2h or 7h of Tg treatment. The heatmaps were ordered according to the order of the clustergram in Fig. 2.

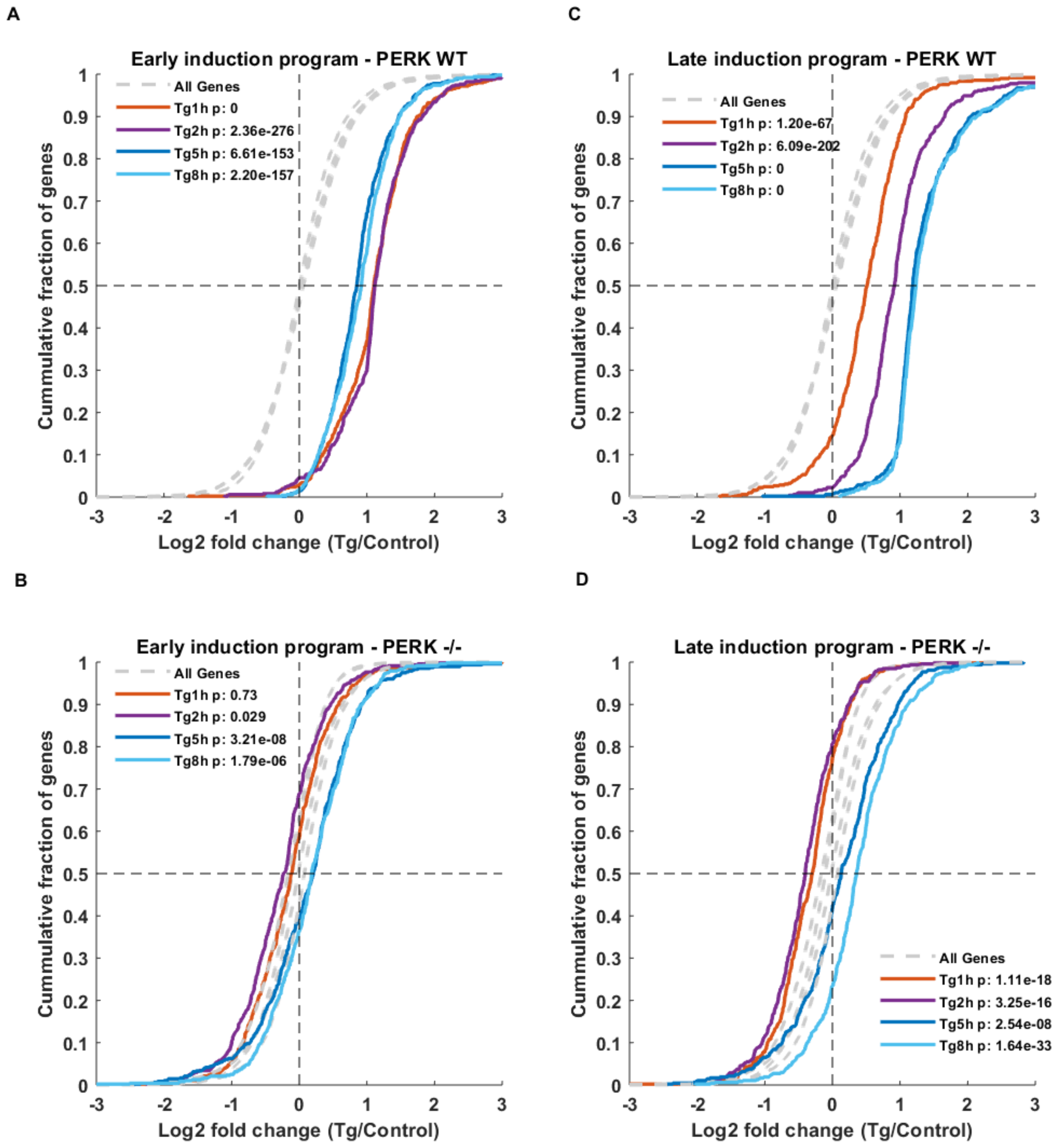


**Figure S5 – Early and late induction gene expression programs are PERK-dependent.** (A,B) CDF plots demonstrate the cumulative fraction of early induction cluster genes (y-axis), as a function of their log2 fold change (LFC) (x-axis) relative to control in (A) PERK WT and (B) PERK -/- cells. Dashed grey lines depict the background LFC distributions of all expressed genes in each timepoint, t-test p-value are indicated. Similar CDF plot analysis was performed for the Late induction cluster (C,D).

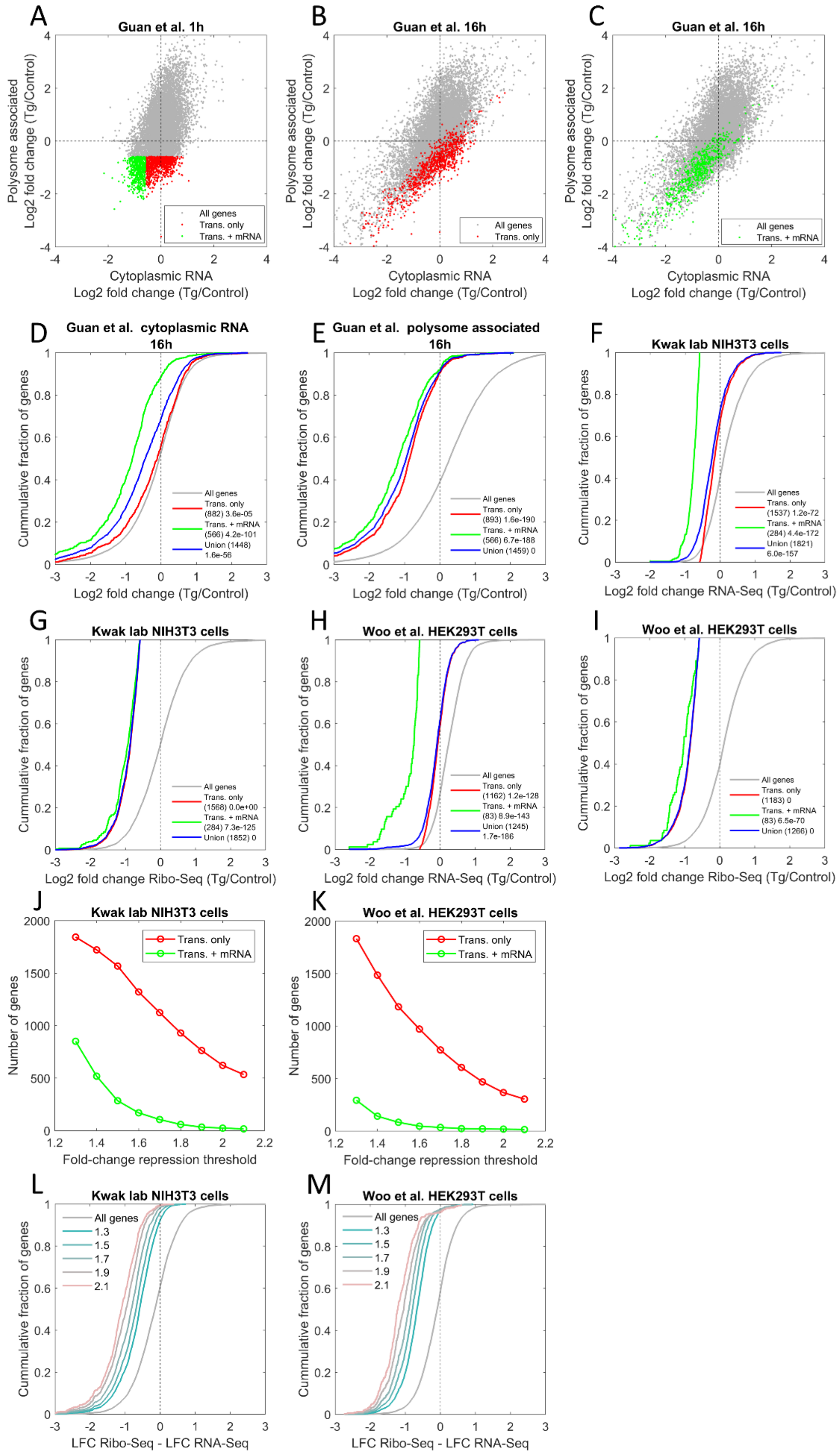

A
Guan et al. 1h
Polysome associated
Log2 fold change (Tg/Control)
Cytoplasmic RNA
Log2 fold change (Tg/Control)
All genes
Trans. only
Trans. + mRNA
B
Guan et al. 16h
Polysome associated
Log2 fold change (Tg/Control)
Cytoplasmic RNA
Log2 fold change (Tg/Control)
All genes
Trans. only
C
Guan et al. 16h
Polysome associated
Log2 fold change (Tg/Control)
Cytoplasmic RNA
Log2 fold change (Tg/Control)
All genes
Trans. + mRNA
D
Guan et al. cytoplasmic RNA
16h
Cummulative fraction of genes
Log2 fold change (Tg/Control)
All genes
Trans. only
(882) 3.6e-05
Trans. + mRNA
(566) 4.2e-101
Union (1448)
1.6e-56
E
Guan et al. polysome associated
16h
Cummulative fraction of genes
Log2 fold change (Tg/Control)
All genes
Trans. only
(893) 1.6e-190
Trans. + mRNA
(566) 6.7e-188
Union (1459) 0
F
Kwak lab NIH3T3 cells
Cummulative fraction of genes
Log2 fold change RNA-Seq (Tg/Control)
All genes
Trans. only
(1537) 1.2e-72
Trans. + mRNA
(284) 4.4e-172
Union (1821)
6.0e-157
G
Kwak lab NIH3T3 cells
Cummulative fraction of genes
Log2 fold change Ribo-Seq (Tg/Control)
All genes
Trans. only
(1568) 0.0e+00
Trans. + mRNA
(284) 7.3e-125
Union (1852) 0
H
Woo et al. HEK293T cells
Cummulative fraction of genes
Log2 fold change RNA-Seq (Tg/Control)
All genes
Trans. only
(1162) 1.2e-128
Trans. + mRNA
(83) 8.9e-143
Union (1245)
1.7e-186
I
Woo et al. HEK293T cells
Cummulative fraction of genes
Log2 fold change Ribo-Seq (Tg/Control)
All genes
Trans. only
(1183) 0
Trans. + mRNA
(83) 6.5e-70
Union (1266) 0
J
Kwak lab NIH3T3 cells
Number of genes
Fold-change repression threshold
Trans. only
Trans. + mRNA
K
Woo et al. HEK293T cells
Number of genes
Fold-change repression threshold
Trans. only
Trans. + mRNA
L
Kwak lab NIH3T3 cells
Cummulative fraction of genes
LFC Ribo-Seq - LFC RNA-Seq
All genes
1.3
1.5
1.7
1.9
2.1
M
Woo et al. HEK293T cells
Cummulative fraction of genes
LFC Ribo-Seq - LFC RNA-Seq
All genes
1.3
1.5
1.7
1.9
2.1

**Figure S6 – Multiple datasets demonstrate a significant translational component in the repression of ER targets during ER stress.**

(A) Log2 fold change (LFC, 1h Tg/Control) of gene expression in translation (polysome-associated, y axis) vs. mRNA (cytoplasmic RNA, x axis) from Guan et al.[2]. Genes that show repression at 1h of Tg, both at the mRNA and the translational level by 1.5 fold or more are highlighted in green. Genes whose repression was mainly at the level of translation (by 1.5 fold or more) are highlighted in red. (B,C) LFC (16h Tg/Control) of gene expression in translation (y axis) vs. mRNA (x axis) from Guan et al.[2]. Genes whose repression was mainly at the level of translation at 1h of Tg, as defined in (A) (highlighted in red), and genes that were repressed both at the mRNA and the translational levels at 1h of Tg, as defined in (A) (highlighted in green), show similar mode of regulation at 16h. (D,E) CDF plots demonstrate the cumulative fraction of gene groups defined in (A) (y-axis), as a function of their LFC (x-axis) of mRNA (D) or translation (E) at 16h Tg relative to control, largely recapitulate their mode of regulation at 1h. (F, G) Analysis of data from NIH3T3 cells, treated with Tg for 1.5h (Kwak lab GSE103667). CDF plots demonstrate the cumulative fraction of genes for groups defined as repressed both at the mRNA (RNA-seq) and translation (Ribo-seq) levels vs. translation level only (in the same way as in (A)), as a function of their LFC (Tg/Control, x-axis) for RNA-Seq (F) and Ribo-Seq (G). (H, I) Analysis of data from HEK293T cells treated with Tg for 2h (Woo et al.[3]). CDF plots demonstrate the cumulative fraction of genes for groups defined as repressed both at the mRNA and translation levels vs. translation level only (in the same way as in (A)), as a function of their LFC (Tg/Control, x-axis) for RNA-Seq (H) and Ribo-Seq (I). (J-M) Repression analysis for the two additional datasets (from F-I) shows robustness to parameter threshold choice. Similar analysis of separation to mRNA- and translation-level vs. translation-level regulated genes was repeated using multiple cutoff definitions for repression. All cutoffs recapitulated the same trends observed using the 1.5 cutoff (used for F-I and Fig. 5D,E); (J, K) The number of genes in the two groups (repression in both mRNA and translation – green; repression mainly in translation - red) as a function of fold-change repression threshold for the data from NIH3T3 cells (Kwak lab, GSE103667, J) and from HEK293T cells (Woo et al.[3], K). (L, M) CDF plots for the difference between translation log2 fold change (Ribo-Seq LFC) and mRNA-level log2 fold change (RNA-Seq LFC) under multiple repression thresholds in the data from panel J and K respectively, similarly to Fig. 5D,E. In both datasets and under all thresholds, genes were mainly translationally repressed.

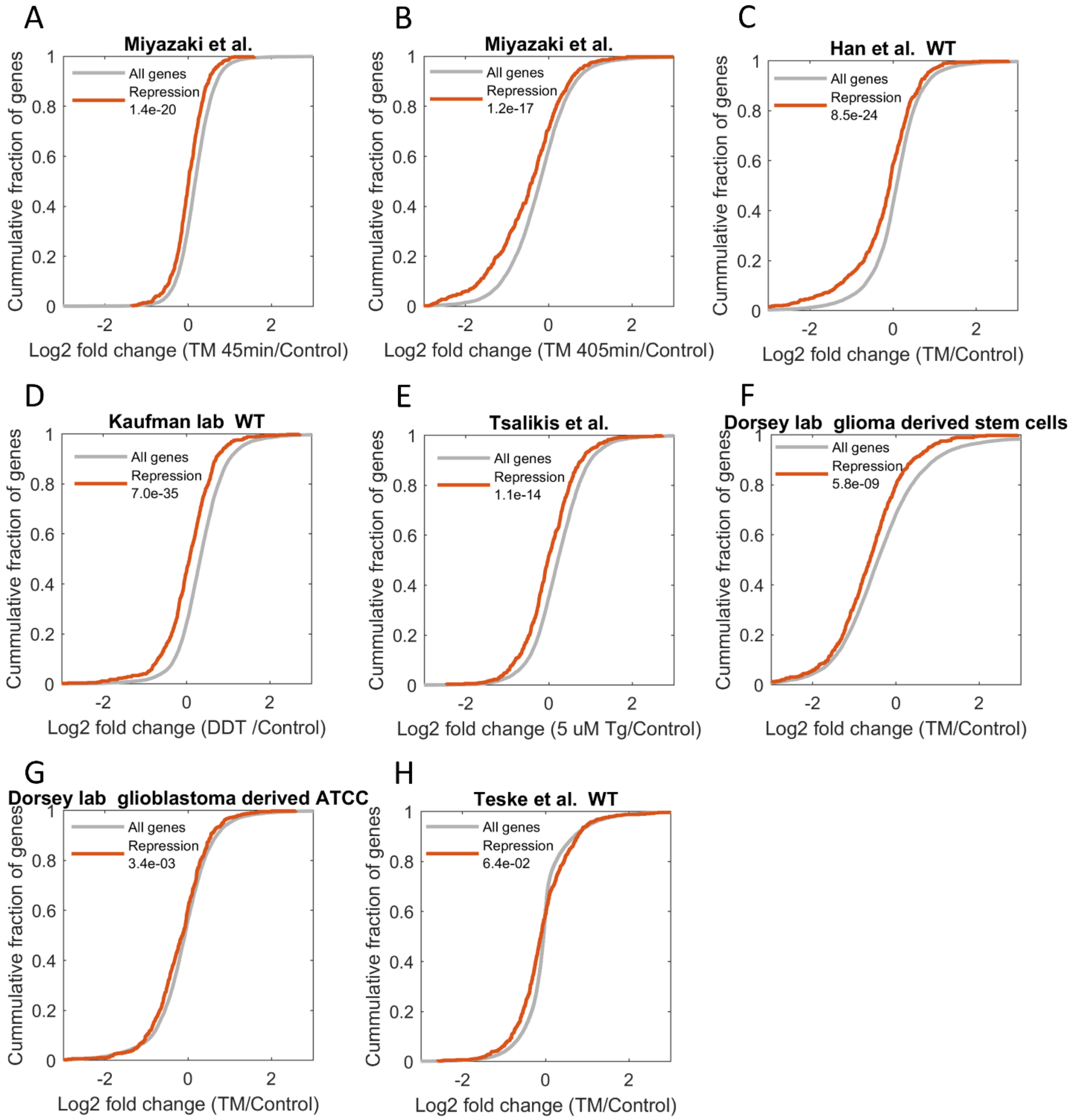


**Figure S7 - Repression program is recapitulated in many cell types and stress treatments.** CDF plots demonstrate the cumulative fraction of each set of genes (y-axis), as a function of their log2 fold change (LFC) (x-axis) relative to control. Background distributions (LFC values of all expressed genes) are marked by grey lines. T-test p-values between each distribution and its respective background are indicated. The analysis demonstrated that the ER stress repression gene expression program that was identified in MEFs (Fig. 2) was recapitulated in many other stress treatments (Tunicamycin, TM, A-C, and DTT, D). Additionally, the ER stress repression gene expression program (Fig. 2) was also recapitulated in other cell types (E-H). See Table S4 for a detailed list of datasets used.

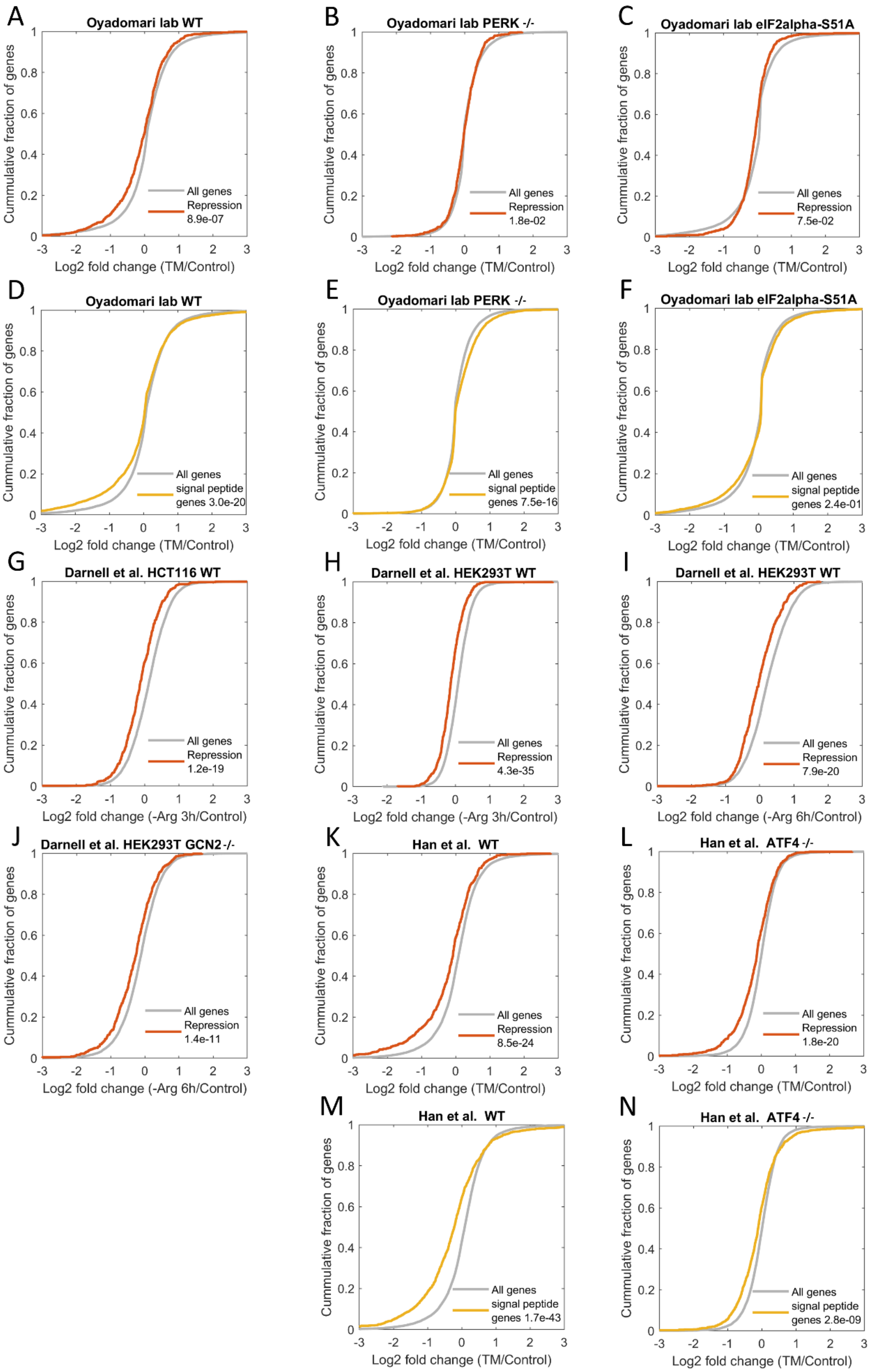
A
Oyadomari lab WT
Cummulative fraction of genes
All genes
Repression
8.9e-07
Log2 fold change (TM/Control)
B
Oyadomari lab PERK -/-
All genes
Repression
1.8e-02
Log2 fold change (TM/Control)
C
Oyadomari lab eIF2alpha-S51A
All genes
Repression
7.5e-02
Log2 fold change (TM/Control)
D
Oyadomari lab WT
All genes
signal peptide
genes 3.0e-20
Log2 fold change (TM/Control)
E
Oyadomari lab PERK -/-
All genes
signal peptide
genes 7.5e-16
Log2 fold change (TM/Control)
F
Oyadomari lab eIF2alpha-S51A
All genes
signal peptide
genes 2.4e-01
Log2 fold change (TM/Control)
G
Darnell et al. HCT116 WT
All genes
Repression
1.2e-19
Log2 fold change (-Arg 3h/Control)
H
Darnell et al. HEK293T WT
All genes
Repression
4.3e-35
Log2 fold change (-Arg 3h/Control)
I
Darnell et al. HEK293T WT
All genes
Repression
7.9e-20
Log2 fold change (-Arg 6h/Control)
J
Darnell et al. HEK293T GCN2-/-
All genes
Repression
1.4e-11
Log2 fold change (-Arg 6h/Control)
K
Han et al. WT
All genes
Repression
8.5e-24
Log2 fold change (TM/Control)
L
Han et al. ATF4 -/-
All genes
Repression
1.8e-20
Log2 fold change (TM/Control)
M
Han et al. WT
All genes
signal peptide
genes 1.7e-43
Log2 fold change (TM/Control)
N
Han et al. ATF4 -/-
All genes
signal peptide
genes 2.8e-09
Log2 fold change (TM/Control)

**Figure S8 - PERK-mediated repression of ER targets during ER stress is largely mediated through eIF2α phosphorylation, and partly dependent on ATF4.**

CDF plots demonstrate the cumulative fraction of each set of genes (y-axis), as a function of their log2 fold change (LFC) (x-axis) relative to control. Background distributions (LFC values of all expressed genes) are marked by grey lines. T-test p-values between each distribution and its respective background are indicated. The analysis examines the potential role of regulators downstream to PERK in the mediation of ER target repression. The ER stress repression program, as well as ER targets (signal peptide containing proteins) showed significant repression in WT MEFs (A and D respectively), which was completely abrogated in PERK -/- MEFs (B,E) and eIF2α-S51A MEFs (C,F), indicating that eIF2α phosphorylation downstream of PERK plays a major role in the preferential repression of ER targets. (G-J). Significant downregulation of the ER stress repression program is shown in HCT116 WT (G), HEK293T WT (H,I), as well as GCN2 knockout HEK293T cells (J) upon Arginine deprivation[4]. See Table S4 for a detailed list of datasets used. (K,L) Significant downregulation in the repression program is observed in WT and ATF4 -/- MEFs[5]. (M,N) ER targets (signal peptide containing proteins) showed significant repression in WT MEFs (M), which is partly alleviated in ATF4 -/- MEFs (N).

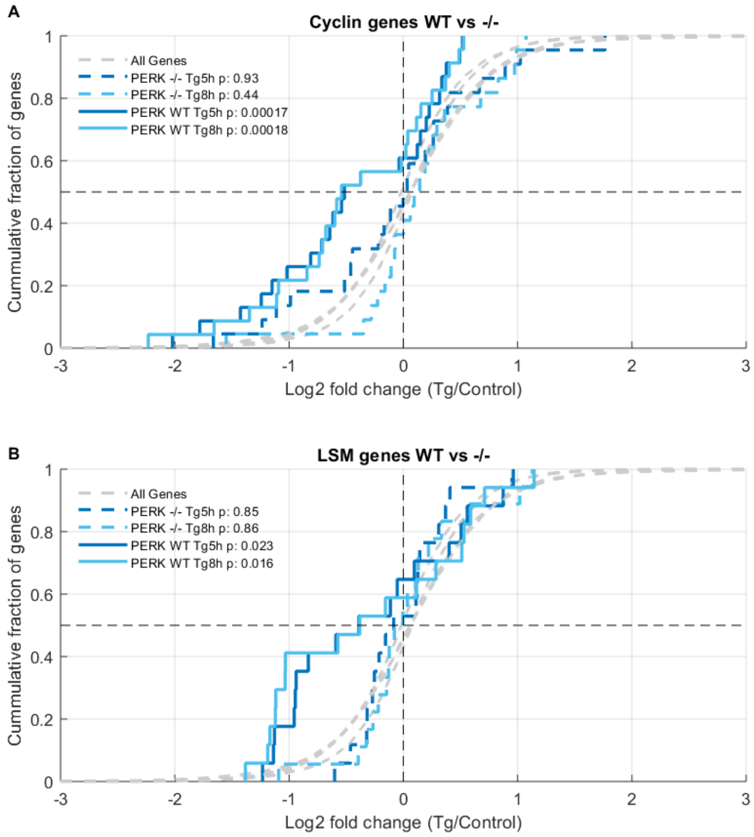


**Figure S9 - Cyclins and LSM domain proteins show a PERK-dependent repression in chronic ER stress.**

CDF plots depict cyclins (A) and LSM-domain proteins (B) log2 fold changes (LFCs) at the 5h and 8h Tg treatments compared to control in either PERK WT (solid lines) or PERK -/- cells (dashed lines), demonstrating a significant PERK-dependent repression of these two sets of genes. Grey dashed lines indicate the background distributions of the LFCs of all expressed genes in the different timepoints, t-test p-values between each distribution and its respective background are indicated.

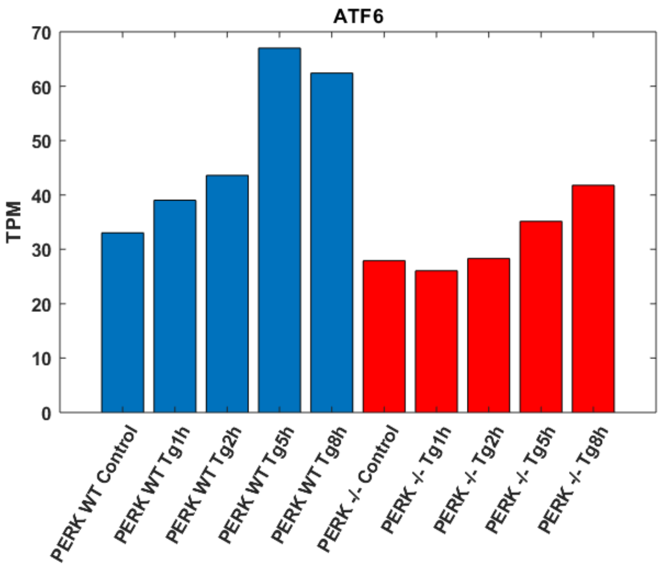


**Figure S10 – ATF6 induction in ER stress is partially PERK-dependent.**
TPM expression levels of ATF6 are shown in the different samples.

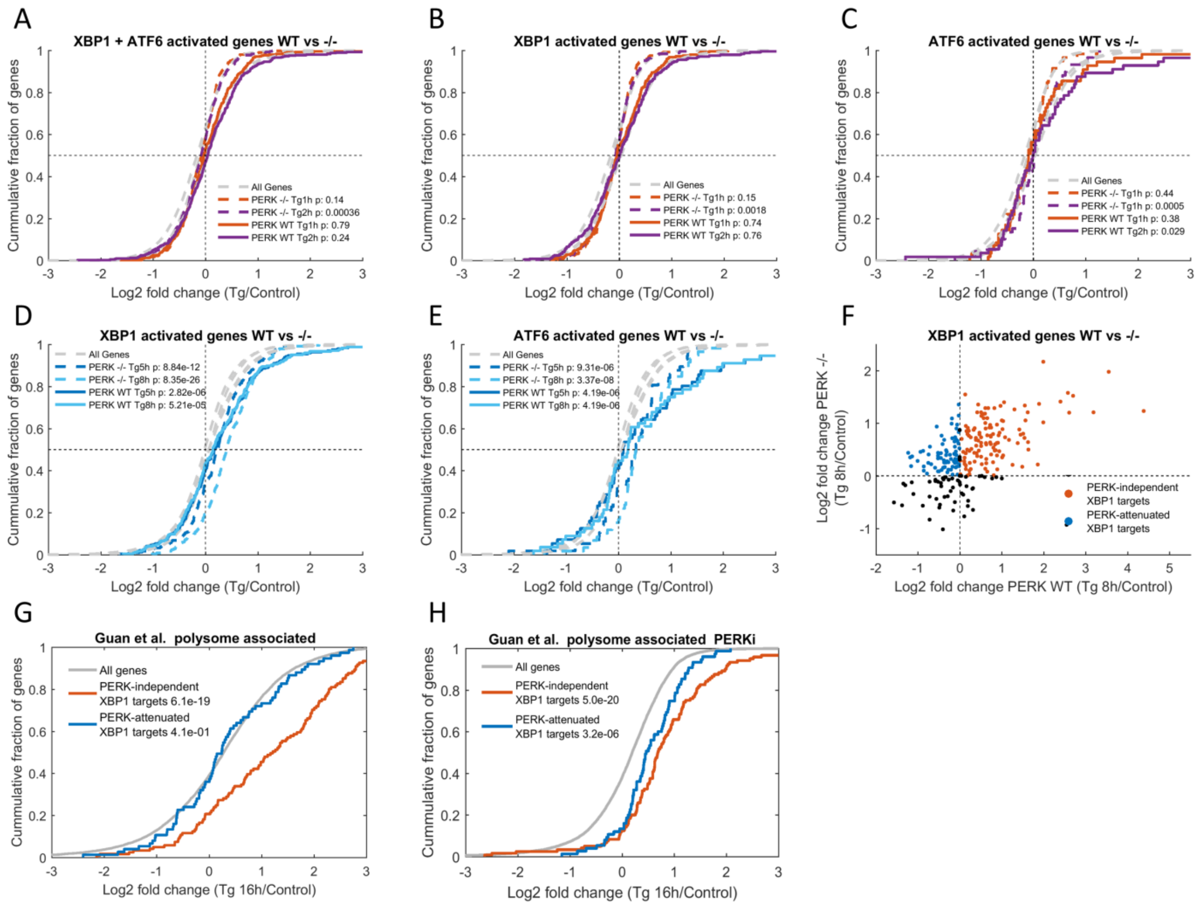


**Figure S11 – Complex interplay between XBP1 and ATF6 target genes and PERK.**

(A) Marginal change in XBP1-ATF6 targets (defined by Shoulders et al.[6]) levels in the early timepoints of ER stress as demonstrated by CDF plots of the log2 fold changes (LFCs) at 1h or 2h post Tg vs. control. (B) the same trend holds for XBP1 targets and (C) ATF6 targets separately. (D-E) The same analysis as in Fig. 6A, B holds true for XBP1 targets. (F) Definition of XBP1 independent (up-regulated in PERK WT and -/-, red) and attenuated (up-regulated only in PERK -/-, blue) groups. (G,H) The same analysis as in Figure 6C,D holds true for XBP1 targets. CDF plot shows that PERK-independent XBP1 targets are induced in the Guan et al. dataset after 16h of Tg treatment (G, red curve) irrespective of PERK inhibition (H, red curve), while PERK-attenuated XBP1 targets were unchanged at 16h Tg treatment (G, blue curve), and induced with the inhibition of PERK (H, blue curve). ATF6 target set subsets were too small to draw conclusions. Grey lines indicate the background distributions of the LFCs of all expressed genes in the different conditions, t-test p-values between each distribution and its respective background are indicated.

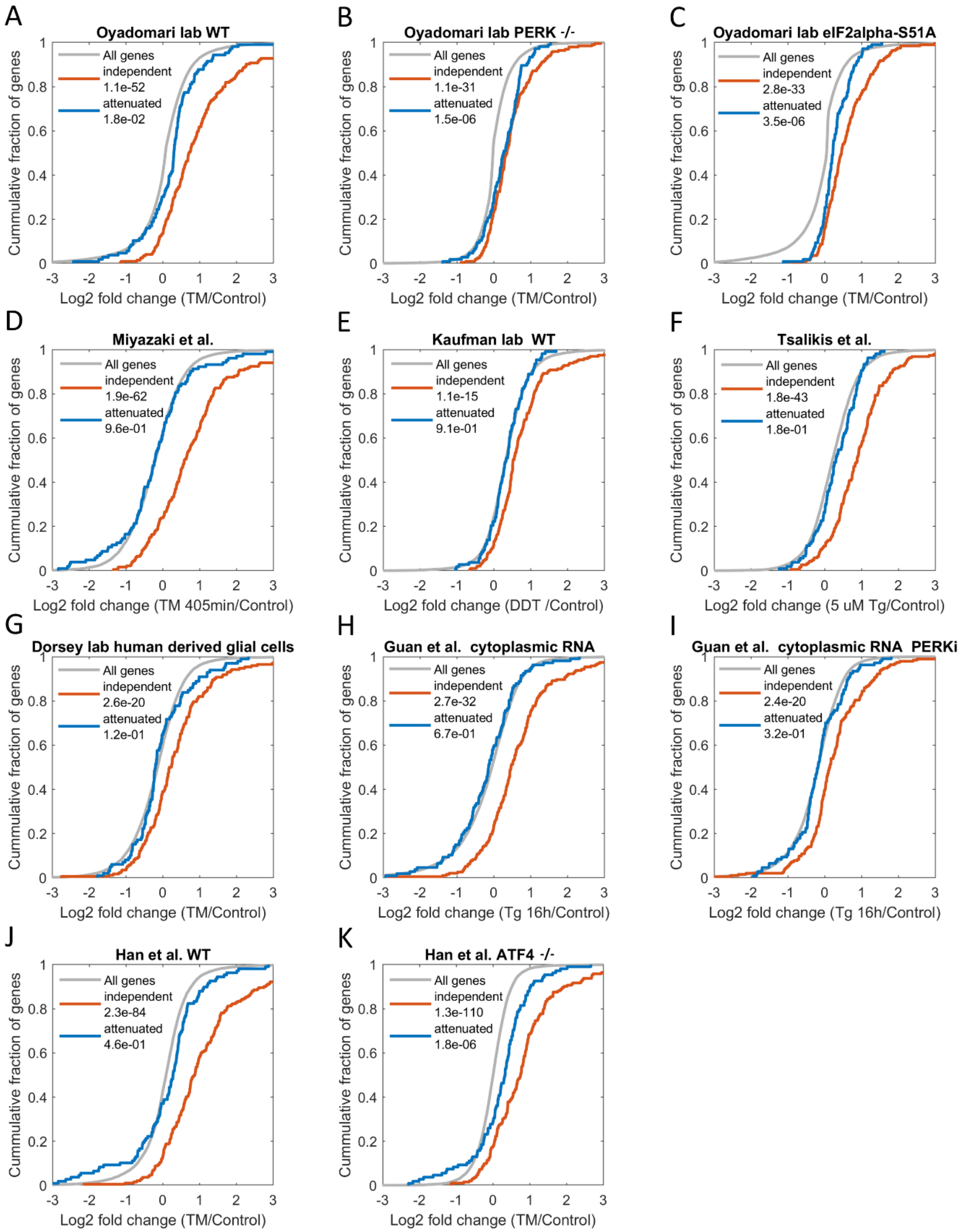
A
Oyadomari lab WT
All genes
independent
1.1e-52
attenuated
1.8e-02
Cummulative fraction of genes
Log2 fold change (TM/Control)
B
Oyadomari lab PERK -/-
All genes
independent
1.1e-31
attenuated
1.5e-06
Cummulative fraction of genes
Log2 fold change (TM/Control)
C
Oyadomari lab eIF2alpha-S51A
All genes
independent
2.8e-33
attenuated
3.5e-06
Cummulative fraction of genes
Log2 fold change (TM/Control)
D
Miyazaki et al.
All genes
independent
1.9e-62
attenuated
9.6e-01
Cummulative fraction of genes
Log2 fold change (TM 405min/Control)
E
Kaufman lab WT
All genes
independent
1.1e-15
attenuated
9.1e-01
Cummulative fraction of genes
Log2 fold change (DDT /Control)
F
Tsalikis et al.
All genes
independent
1.8e-43
attenuated
1.8e-01
Cummulative fraction of genes
Log2 fold change (5 uM Tg/Control)
G
Dorsey lab human derived glial cells
All genes
independent
2.6e-20
attenuated
1.2e-01
Cummulative fraction of genes
Log2 fold change (TM/Control)
H
Guan et al. cytoplasmic RNA
All genes
independent
2.7e-32
attenuated
6.7e-01
Cummulative fraction of genes
Log2 fold change (Tg 16h/Control)
I
Guan et al. cytoplasmic RNA PERKi
All genes
independent
2.4e-20
attenuated
3.2e-01
Cummulative fraction of genes
Log2 fold change (Tg 16h/Control)
J
Han et al. WT
All genes
independent
2.3e-84
attenuated
4.6e-01
Cummulative fraction of genes
Log2 fold change (TM/Control)
K
Han et al. ATF4 -/-
All genes
independent
1.3e-110
attenuated
1.8e-06
Cummulative fraction of genes
Log2 fold change (TM/Control)

**Figure S12 – PERK-mediated attenuation of XBP1-ATF6 targets is governed by transcriptional and translational regulation, and involves eIF2α phosphorylation and ATF4.**

Changes in the levels of PERK-independent XBP1-ATF6 targets (red curves) and PERK-attenuated XBP1-ATF6 targets (blue curves) (XBP1-ATF6 targets taken from Shoulders et al.[6], groups defined in Fig. 6B) during ER stress, as demonstrated by CDF plots of the log2 fold changes (LFCs) of treatment vs. control. Grey lines indicate the background distributions of the LFCs of all expressed genes in the different conditions, t-test p-values between each distribution and its respective background are indicated. (A-C) PERK-attenuated XBP1-ATF6 targets were unchanged at TM treatment in PERK WT MEFs (A). Relief in PERK-mediated inhibition of the PERK-attenuated XBP1-ATF6 targets in the PERK -/- MEFs (B) and in eIF2α-S51A MEFs (C). (D-G) Similar trends of induction vs. attenuation for the PERK-independent and PERK-attenuated XBP1-ATF6 targets were found in several mRNA expression datasets, including response to TM[7] (D) and DTT (E), as well as in other cell types, including Tg treated intestinal stem cells[8] (F), and glial cells treated with TM (G). (H,I) Analysis of mRNA expression from Guan et al.[2] showed that, unlike the translational level (Fig. 6C,D), no relief was observed at the mRNA level of the PERK-attenuated XBP1-ATF6 targets upon PERK inhibition for 4h. (J,K) Significant alleviation of repression for the PERK-attenuated XBP1-ATF6 targets in ATF4 -/- MEFs (K) compared to WT MEFs (J) from Han et al.[5]

See Table S4 for a detailed list of datasets used.

**References:**


1 Shalgi, R. *et al.* Widespread regulation of translation by elongation pausing in heat shock. *Molecular cell* **49**, 439-452, doi:10.1016/j.molcel.2012.11.028 (2013).

2 Guan, B. J. *et al.* A Unique ISR Program Determines Cellular Responses to Chronic Stress. *Mol Cell* **68**, 885-900 e886, doi:10.1016/j.molcel.2017.11.007 (2017).

3 Woo, Y. M. *et al.* TED-Seq Identifies the Dynamics of Poly(A) Length during ER Stress. *Cell Rep* **24**, 3630-3641 e3637, doi:10.1016/j.celrep.2018.08.084 (2018).

4 Darnell, A. M., Subramaniam, A. R. & O'Shea, E. K. Translational Control through Differential Ribosome Pausing during Amino Acid Limitation in Mammalian Cells. *Mol Cell* **71**, 229-243 e211, doi:10.1016/j.molcel.2018.06.041 (2018).

5 Han, J. *et al.* ER-stress-induced transcriptional regulation increases protein synthesis leading to cell death. *Nat Cell Biol* **15**, 481-490, doi:10.1038/ncb2738 (2013).

6 Shoulders, M. D. *et al.* Stress-independent activation of XBP1s and/or ATF6 reveals three functionally diverse ER proteostasis environments. *Cell Rep* **3**, 1279-1292, doi:10.1016/j.celrep.2013.03.024 (2013).

7 Miyazaki, Y., Chen, L. C., Chu, B. W., Swigut, T. & Wandless, T. J. Distinct transcriptional responses elicited by unfolded nuclear or cytoplasmic protein in mammalian cells. *Elife* **4**, doi:10.7554/eLife.07687 (2015).

8 Tsalikis, J. *et al.* The transcriptional and splicing landscape of intestinal organoids undergoing nutrient starvation or endoplasmic reticulum stress. *BMC Genomics* **17**, 680, doi:10.1186/s12864-016-2999-1 (2016).

**Table S4 – List of all datasets used.**

| Dataset | Accession | Assay | Cell type | Genotype | Condition |
|---|---|---|---|---|---|
| Guan et al. Mol Cell 2017[1] | GSE90070 | RNA-Seq of polysome-associated and cytoplasmic RNA | MEF | WT | Tg 1hr<br>Tg 16hr<br>Tg 16hr + PERK inhibitor |
| Woo et al. Cell Reports 2018 [2] | GSE103719 | RNA-Seq + Ribo-Seq | HEK293T cells | WT | Tg 2 hr |
| Kwak lab NIH3T3 cells | GSE103667 | RNA-Seq + Ribo-Seq | NIH3T3 | WT | Tg 1.5 hr |
| Miyazaki et al. Elife 2015 [3] | GSE65636 | RNA-Seq | NIH3T3 | WT | TM 45 min<br>TM 405 min |
| Oyadomari lab | GSE49598 | Microarrays | MEF | WT<br>PERK -/-<br>eIF2α-S51A | TM 12 hr |
| Tsalikis et al. BMC Genomics 2016[4] | GSE84989 | RNA-Seq | Mouse intestinal stem cell | WT | Tg 4 hr |
| Kaufman lab | GSE84450 | RNA-Seq | MEF | WT | DTT 4 hr |
| Dorsey lab | GSE102505 | RNA-Seq | -Human astrocytes<br>-Glioma derived stem cells<br>-human glioblastoma | WT | TM 24 hr |
| Darnell et al. Mol Cell 2018[5] | GSE113751 | Ribo-Seq | HEK293T (WT and GCN2 -/-)<br>HCT116 (WT) | WT<br>GCN2 -/- | Arg deprivation for 3 and 6 hr |
| Teske et al. Mol biology of the cell 2011[6] | GSE29929 | microarrays | mouse liver cells | WT | TM 6 hr |

| | | | | | |
|---|---|---|---|---|---|
| Han et al. Nature cell biology 2013[7] | GSE35681 | RNA-Seq | MEF | WT<br>ATF4 -/- | TM 8 hr |

**References:**


1 Guan, B. J. *et al.* A Unique ISR Program Determines Cellular Responses to Chronic Stress. *Mol Cell* **68**, 885-900 e886, doi:10.1016/j.molcel.2017.11.007 (2017).

2 Woo, Y. M. *et al.* TED-Seq Identifies the Dynamics of Poly(A) Length during ER Stress. *Cell Rep* **24**, 3630-3641 e3637, doi:10.1016/j.celrep.2018.08.084 (2018).

3 Miyazaki, Y., Chen, L. C., Chu, B. W., Swigut, T. & Wandless, T. J. Distinct transcriptional responses elicited by unfolded nuclear or cytoplasmic protein in mammalian cells. *Elife* **4**, doi:10.7554/eLife.07687 (2015).

4 Tsalikis, J. *et al.* The transcriptional and splicing landscape of intestinal organoids undergoing nutrient starvation or endoplasmic reticulum stress. *BMC Genomics* **17**, 680, doi:10.1186/s12864-016-2999-1 (2016).

5 Darnell, A. M., Subramaniam, A. R. & O'Shea, E. K. Translational Control through Differential Ribosome Pausing during Amino Acid Limitation in Mammalian Cells. *Mol Cell* **71**, 229-243 e211, doi:10.1016/j.molcel.2018.06.041 (2018).

6 Teske, B. F. *et al.* The eIF2 kinase PERK and the integrated stress response facilitate activation of ATF6 during endoplasmic reticulum stress. *Mol Biol Cell* **22**, 4390-4405, doi:10.1091/mbc.E11-06-0510 (2011).

7 Han, J. *et al.* ER-stress-induced transcriptional regulation increases protein synthesis leading to cell death. *Nat Cell Biol* **15**, 481-490, doi:10.1038/ncb2738 (2013).
